# TBX3 is essential to establish the posterior boundary of anterior genes and up-regulate posterior genes with HAND2 during onset of limb bud development

**DOI:** 10.1101/2024.01.30.577998

**Authors:** Geoffrey Soussi, Ausra Girdziusaite, Shalu Jhanwar, Victorio Palacio, Rushikesh Sheth, Rolf Zeller, Aimée Zuniga

**Author notes:** these authors contributed equally (Ausra Girdziusaite, Shalu Jhanwar). **Summary statement** It was unknown how anterior gene expression is excluded from the posterior limb bud mesenchyme. We show that TBX3 repression sets the posterior boundaries and promotes posterior identity with HAND2.

## Abstract

During limb bud formation, axes polarities are established as evidenced by the spatially restricted expression of key regulator genes. In particular, the mutually antagonistic interaction between the GLI3 repressor and HAND2 results in the distinct and non-overlapping anterior-distal *Gli3* and posterior *Hand2* expression domains. This hallmarks establishment of antero-posterior (AP) limb axis polarity together with spatially restricted expression of homeodomain and other transcriptional regulators. Here, we identify TBX3 as the transcription factor to initiate AP axis polarity in mouse limb buds. ChIP-seq and differential gene expression analysis of wildtype and mutant limb buds identifies the TBX3-specific and shared TBX3-HAND2 target genes. High sensitivity fluorescent whole mount *in situ* hybridisation shows that the posterior expression boundaries of anterior genes are positioned by TBX3-mediated repression, which excludes anterior genes such as *Gli3*, *Alx4*, *Hand1* and *Irx3/5* from the posterior limb bud mesenchyme. This exclusion delineates the posterior mesenchymal territory competent to establish the *Shh*-expressing limb bud organiser. In turn, HAND2 is required for *Shh* activation and cooperates with TBX3 to up-regulate shared posterior identity target genes in early limb buds.

## INTRODUCTION

The developing vertebrate limb bud is an experimental paradigm to study the fundamental mechanisms, gene regulatory networks (GRNs) and cellular interactions that govern organ development. During limb bud formation, the antero-posterior (AP), proximo-distal (PD) and dorso-ventral (DV) limb bud axes are established, which results in precise positioning of the two main signal centers, the Fibroblast growth factor (FGF) signaling apical ectodermal ridge (AER) and the Sonic hedgehog (SHH)- signaling posterior mesenchymal organizer (reviewed by Zuniga, 2015). Many of the transcriptional regulators and signaling pathways that control the onset of limb bud development have been identified and functionally analyzed (reviewed by Zuniga and Zeller, 2020). One such hallmark is the AP polarization of the mesenchyme during limb bud formation that manifests itself by the complementary expression of *Gli3* in the anterior-distal and *Hand2* in the posterior mesenchyme (Welscher et al., 2002a). These two transcriptional regulators act in mutually antagonistic as genetic inactivation of *Gli3* in mouse limb buds causes anterior expansion of *Hand2* expression and pre-axial (anterior) digit polydactyly (Welscher et al., 2002a; Lopez-Rios et al., 2012). Conversely, *Gli3* expands posteriorly and establishment of posterior limb bud identity and *Shh* activation are disrupted in *Hand2-*deficient limb buds, which phenocopies the digit loss observed in *Shh-*deficient mouse limbs (Galli et al., 2010). Initially, the GLI3 repressor isoform (GLI3R) and HAND2 are co-expressed, but the antagonism results in rapid establishment of their complementary expression domains during limb bud formation (Osterwalder et al., 2014). A recent study has however cast doubt on the postulated repressive function of the predominant GLI3R isoform (Wang et al., 2000) upstream on *Hand2* prior to activation of SHH signaling (Lex et al., 2022). Genetic loss-of-function analysis showed that initially *Hand2* and then SHH signaling are required to overcome the repressive effect of GLI3R the posterior mesenchyme, which is essential for digit patterning and distal progression of limb bud outgrowth (Litingtung et al., 2002; Welscher et al., 2002b; Zhu et al., 2008; Galli et al., 2010). Therefore, an important open question is whether the *Gli3* and *Hand2* expression boundaries are established by direct cross-regulation between HAND2 and GLI3R or depend on unknown additional transcriptional regulators. The present study addresses posterior boundary formation for anterior-proximal and anterior-distal gene expression during establishment of the AP limb bud axis.

Several transcription factors (TFs) required for activation/up-regulation of *Gli3* and *Hand2* have been identified in mouse limb buds. For example, positive regulation of *Gli3* depends on *Irx3/Irx5* and *Sall4* that impact *Gli3* expression via a specific enhancer active in the anterior mesenchyme (Li et al., 2014; Akiyama et al., 2015). In turn, *Gli3* enhances the expression of *Irx3/Irx5* and *Sall4* as part of a transcriptional feedback (Yokoyama et al., 2017). The expression of *Hand2* in the posterior limb bud mesenchyme depends on specific homeodomain TFs. *Hox9* paralogues are required in a redundant fashion to establish the posterior *Hand2* expression domain in early forelimb buds, while *Isl1* acts upstream of *Hand2* in hindlimb buds (Xu and Wellik, 2011; Itou et al., 2012). The *Pbx1* and *Pbx2* homeoproteins function in up-regulation of *Hand2* expression by interacting with specific enhancers (Capellini et al., 2006; Losa et al., 2023). In the posterior limb bud mesenchyme, PBX1 forms a complex with HAND2 to co-regulate target genes that include several key regulators of mouse limb bud development (Losa et al., 2023). Together, these studies reveal the complexity of gene regulatory interactions among different key players, but fall short of providing insights into the repressive mechanisms that restrict *Gli3* and *Hand2* expression during AP axis polarisation. (Osterwalder et al., 2014) identified the HAND2 target *cis*-regulatory modules (CRMs) in the genomic landscapes of genes functioning during the onset of limb bud development. This analysis identified an enhancer in the *Gli3* genomic landscape, which suggested that HAND2 might directly repress *Gli3* in the posterior limb bud mesenchyme. Moreover, *Tbx3* was identified as a HAND2 target gene in posterior limb bud mesenchyme that is in a complementary manner to *Gli3*. Together with analysis of mutant limb buds, this led to the proposal that HAND2 might repress *Gli3* in the posterior mesenchyme and that *Tbx3* might participate in “fine-tuning” the posterior *Gli3* boundary (Osterwalder et al., 2014).

*Tbx3* is a member of the *Tbx2* sub-family of T-box TFs (reviewed by Sheeba and Logan, 2017) and is first expressed in the lateral plate mesenchyme, then in the posterior and anterior mesenchyme and AER during early limb bud development (Emechebe et al., 2016). Haploinsufficient *TBX3* mutations in humans cause the ulnar-mammary syndrome, which results in a pleiotropic phenotype that includes severe reductions of posterior skeletal elements of the upper extremities (Bamshad et al., 1999). Analysis of *Tbx3*-deficient mouse embryos shows that TBX3 acts initially upstream of *Hand2*, but detailed analysis was precluded due to the mid-gestational lethality (Davenport et al., 2003; Frank et al., 2013). Conditional inactivation of *Tbx3* in early mouse limb buds circumvents this lethality and shows that *Tbx3* is required in the posterior mesenchyme to up-regulate *Hand2* and *Shh* expression (Emechebe et al., 2016). In turn, the upregulation and maintenance of posterior *Tbx3* expression depends on both *Hand2* and SHH signaling (Galli et al., 2010). In the anterior limb bud mesenchyme, *Tbx3* expression is activated slightly later and requires SMAD4-mediated BMP signal transduction (Emechebe et al., 2016; Gamart et al., 2021). The anterior TBX3 protein colocalises with the GLI3 protein to primary cilia and interacts with the SUFU/Kif7 complex to stabilize the GLI3 full-length (GLI3FL) protein and its processing to the GLI3R isoform (Emechebe et al., 2016). In *Tbx3*-deficient limb buds, both GLI3 protein isoforms are degraded, which results in a characteristic pre-axial polydactyly with similarities to *Gli3*-deficient forelimb buds (Lopez-Rios et al., 2012; Emechebe et al., 2016). The knowledge gained from studying the early gene regulatory interactions that control AP limb axis formation and the similarities of mouse limb skeletal phenotypes with human congenital limb malformations provides important insights into their aetiology and is relevant to identification of human disease alleles (Tao et al., 2017).

For this study, we have generated a novel mouse *Tbx3* allele with a 3xFLAG epitope tag inserted into the carboxy-terminal part of the TBX3 protein. This *Tbx3*^3xF^ allele was used for ChIP-seq and combined with open chromatin analysis (ATAC-seq) and differential gene expression analysis (RNA-seq) from wildtype and mutant forelimb buds to identify the TBX3 cistrome and target genes during onset of limb bud development. In parallel, HAND2 target genes were used to identify the target genes shared betweenTBX3 and HAND2. Spatial analysis of target gene expression and enhancer activities in wildtype and mutant limb buds resulted in several major findings. *Tbx3* expression in the anterior mesenchyme is upregulated in *Hand2*-deficient limb buds, while its expression in the posterior mesenchyme depends critically on *Hand2*. Fluorescent HCR^TM^ RNA *in situ* analysis reveals that *Tbx3* is required on its own for establishing the precise posterior expression boundaries of anterior genes such as *Gli3*, *Hand1*, *Alx4* and *Irx3/5*. This finding is important in light of previous studies showing that *Hand2* and *Shh* need to override the repressive activity of GLI3R to enable progression of limb bud development (Litingtung et al., 2002; Welscher et al., 2002b; Galli et al., 2010). Analysis of the two CRMs that regulate the spatial *Gli3* expression in mouse limb buds (Osterwalder et al., 2018), establishes that TBX3 interacts and restricts their activity to the posterior boundary of the *Gli3* expression domain. This excludes *Gli3* enhancer activity and transcription from the posterior mesenchymal territory in which *Shh*-expression is activated. This study identifies an unexpected unique function for TBX3 in positioning AP expression boundaries, which is a crucial step in enabling establishment of the SHH signaling center in the posterior limb bud mesenchyme. In the posterior mesenchyme, *Tbx3* functions together with *Hand2* in positive regulation of posterior identity and *Shh* pathway genes.

## RESULTS

### Identification of TBX3 target genes in mouse forelimb buds

A 3xFLAG (3xF) epitope tag was inserted by homologous recombination into the carboxy-terminal part of the mouse *Tbx3* open reading frame. This *Tbx3*^3xF^ allele allows specific detection of endogenous TBX3^3xF^ proteins using anti-FLAG M2 antibodies in mouse wildtype embryos and forelimb buds (Fig. 1A, 1B and Fig. S1A-C). The insertion of the 3xFLAG epitope tag does not alter the spatial distribution of *Tbx3*^3xF^ transcripts (Fig. 1C) and the TBX3^3xF^ protein is predominantly nuclear in the posterior and anterior limb bud domains (E10.5, 33-35 somites, Fig. 1D-F). *Tbx3*^3xF/3xF^ mice are born at the expected Mendelian ratios and display no overt limb or other phenotypes. Forelimb buds of *Tbx3*^3xF/3xF^ embryos were used to identify the TBX3 cistrome in early mouse forelimb buds by chromatin immunoprecipitation in combination with next generation sequencing (ChIP-seq, Fig. 2A and Fig. S1D-F). Two biological replicates consisting each of ∼70 dissected forelimb buds at mouse embryonic day E9.75-E10.25 (29 to 32 somites) were analysed. Statistical analysis of the two replicates by MACS and MSPC identified a common set of 11’422 TBX3^3xF^ ChIP-seq peaks. Roughly equal fractions the TBX3^3xF^ ChIP-seq peaks are located either close to the transcriptional start sites or at a distance of more than 100kb and the peak summits are conserved more than flanking regions (Fig. S1D-F and Table S1). Next, the TBX3 peaks located in regions of accessible/open chromatin were identified by overlapping the ChIP-seq dataset with an ATAC-seq dataset from wildtype forelimb buds at E9.75 (28-29 somites). This identifies 3057 TBX3 interacting regions located in open chromatin, which is a hallmark of promoters and CRMs/enhancers (Fig. 2A, Table S2). HOMER de novo and known motif enrichment analysis of these 3057 regions (Fig. 2B, Fig. S2A) identifies a variety of enriched transcription factor (TF) binding motifs in early mouse limb buds (∼E9.75) that include the motif for the Eomes T-box TF (Fig. 2B). The core region of the *Eomes* binding motif is very similar to if not identical to other *Tbx* motifs such as the conserved core of *Tbx3* motif (Fig. S2B). Furthermore, the genomic regions enriched in TBX3 chromatin complexes also encode binding motifs for other developmental TFs such as zinc finger and homeodomain proteins, which points to possible coregulation by other TF families (Fig. 2B).

**Figure 1.**
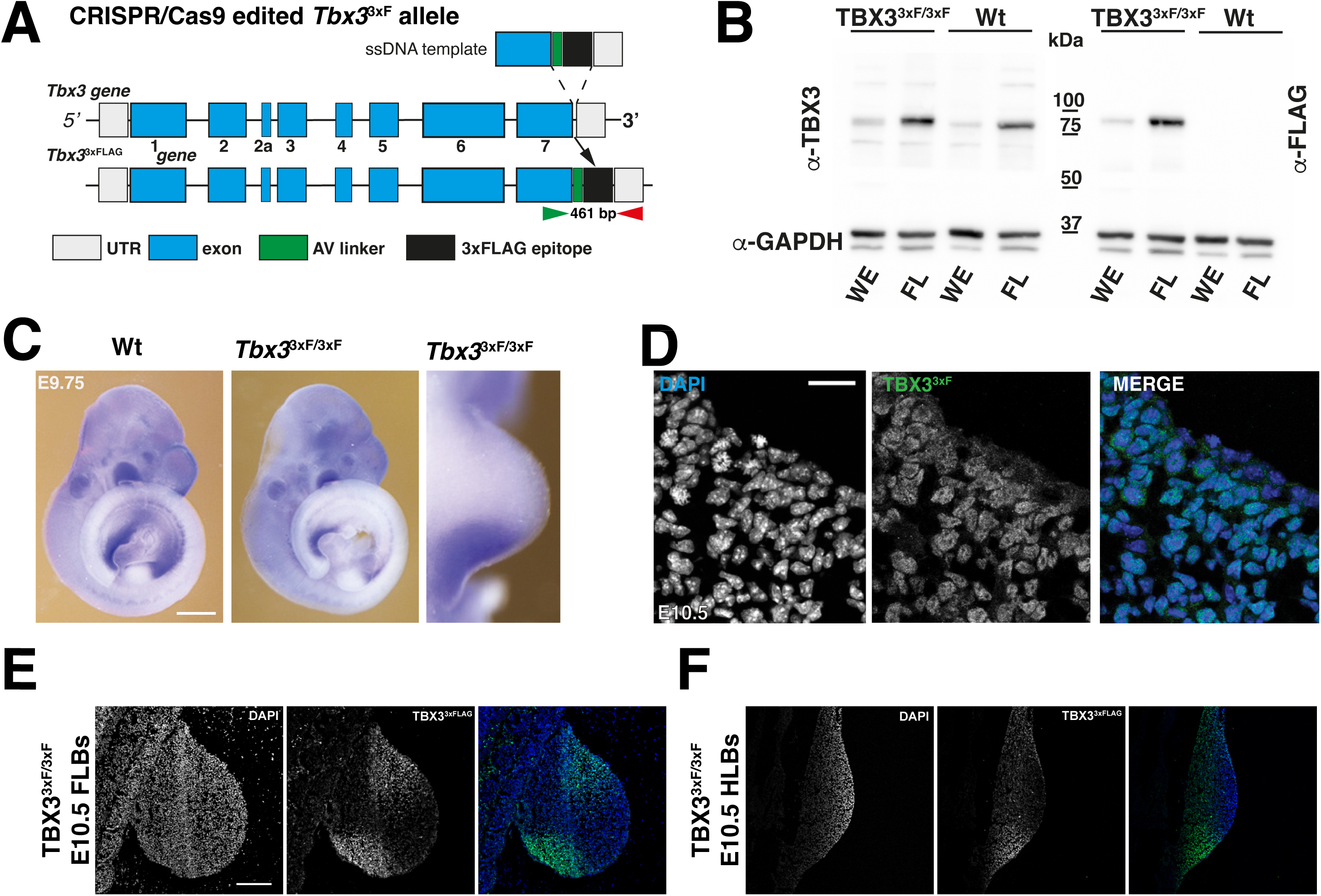
Generation and characterisation of the *Tbx3*^3xFLAG^ (*Tbx3*^3xF^)tagged mouse allele. (A) Scheme summarizing the genome editing process to generate the in-frame 3xFLAG tag epitope insertion (75bp in-frame insertion) into the last *Tbx3* coding exon. The green and red arrowheads indicate the primers for genotyping. (B) Western blot analysis of protein extracts prepared from whole embryos (WE) and forelimb buds (FL) at E10.5. While the TBX3 antibody detects both the wildtype (Wt) and 3xFLAG tagged protein (TBX3^3xF/3xF^,left panel), the M2 FLAG antibodies specifically detect TBX3^3xFLAG^ protein (right panel, n=3). (C) Comparative WISH analysis of Wt and *Tbx3*^3xF/3xF^ mouse embryo at E9.75 (28-30 somites) shows that the spatial *Tbx3* expression is not altered (n=3). Scale bar: 250µm. (D-F) Immunofluorescence analysis of limb bud sections at E10.5 (34-36 somites) shows the nuclear localization of the TBX3^3xF^ protein (panel D). Scale bar: 20µm. It also reveals that the spatial distribution of TBX3 in fore- and hindlimb bud sections correlates well with the known distribution of the wildtype TBX3 protein. All forelimb buds are oriented with anterior to the top and posterior to the bottom in panels C-E. Scale bar: 200µm.

**Figure 2.**
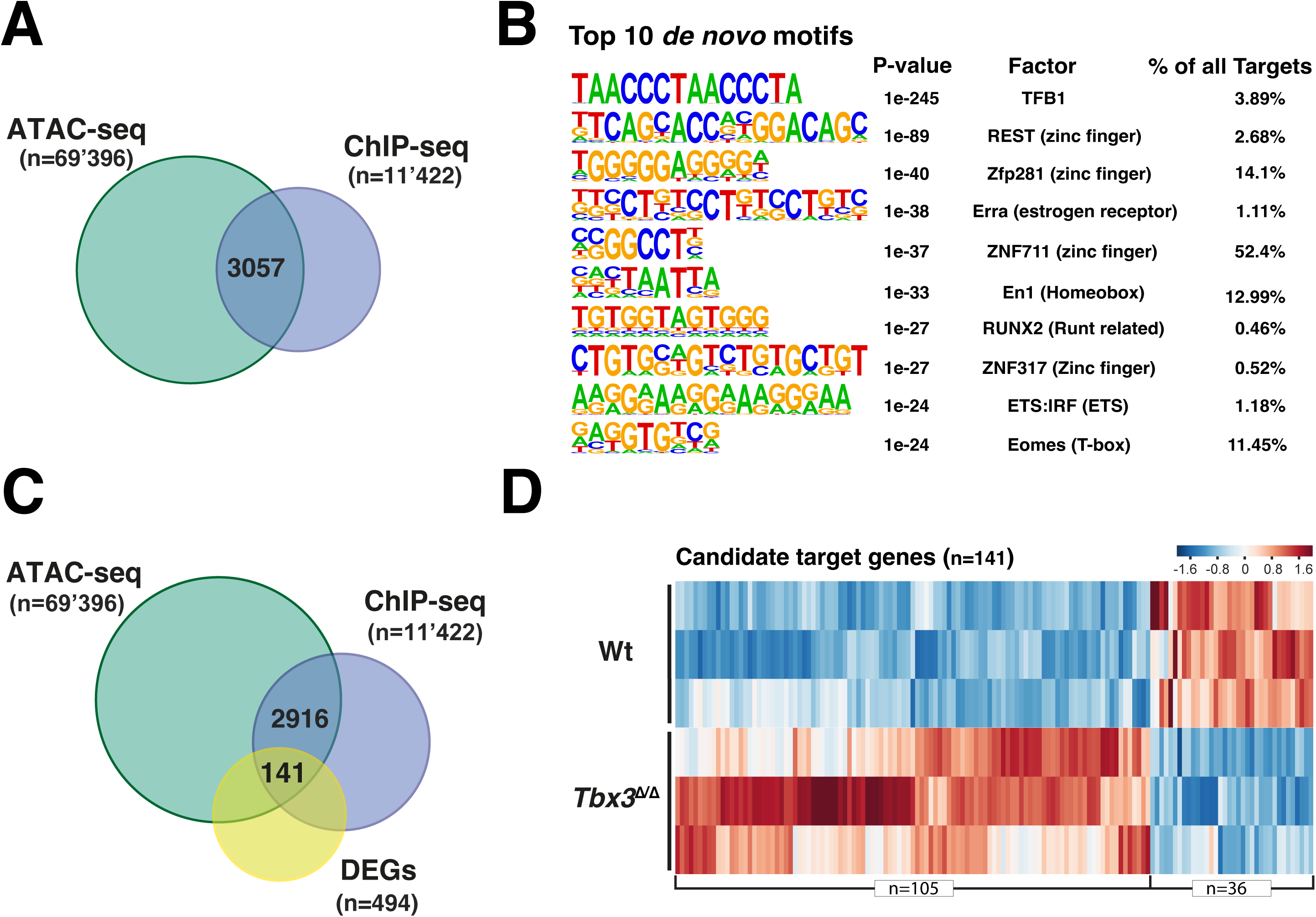
Identification of the TBX3^3xF^ target genes in early mouse forelimb buds. (A) Overlapping regions of open chromatin (ATAC-seq) with the TBX3^3xF^ ChIP-seq peaks identifies 3057 regions that may encode potential *cis*-regulatory modules (CRMs). (B) Top 10 de novo motifs analysis of the accessible genomic regions enriched in TBX3 chromatin complexes identifies several transcription factor motifs including a EOMES related T-box motif. (C) The intersection between the TBX3-bound regions (E9.75-E10.25), open chromatin identified by ATAC-seq (E9.75) and differentially expressed genes (E9.75-10.0) comparing wildtype and *Tbx3*-deficient forelimb buds. This analysis identifies 141 TBX3 candidate target genes. (D) Heatmap showing the relative expression of these candidate targets genes in wildtype and *Tbx3*-deficient forelimb buds. The z-score scale represents mean-subtracted regularized log-transformed read counts.

The differentially expressed genes (DEGs) between wildtype and *Tbx3*-deficient mouse forelimb buds (homozygous for the *Tbx3*^ΔVenus^ allele, Kunasegaran et al., 2014) were identified as follows (Fig. S2C). RNA-seq analysis of three independent biological replicates of forelimb bud pairs from wildtype and *Tbx3*^Δ/Δ^ embryos (E9.75-10.0, 28-31 somites) identified a common set of 494 DEGs with a fold change >1.2 (p-value ≤0.05, Table S3). Among the 494 DEGs in*Tbx3*-deficient mouse forelimb buds, 357 are up- and 137 downregulated (Fig. S2C). The TBX3 target genes in early mouse forelimb buds were identified by mapping the ChIP-seq peaks located within accessible chromatin regions to the nearest gene(s) located within ≤1MB interval using GREAT analysis (Fig. 2C, 2D). Among these, differential expression identifies 141 TBX3-dependent candidate target genes in early mouse forelimb buds at E9.75-E10.0 (Fig. 2D, Table S4). The majority of the TBX3 candidate targets genes are upregulated in *Tbx3*-deficient mouse forelimb buds (105 of 141, Fig. 2D), while the remainder are downregulated (36 of 141, Fig. 2D). Gene Ontology (GO) analysis reveals the predominance of genes with functions in embryonic and limb bud development (Fig. S2D, S2E). Forty-one of the 141 TBX3 target genes (28%) are genes coregulated by SHH signaling during subsequent limb bud outgrowth (E10.5, Table S6, Probst et al., 2011). Together, these results indicate that TBX3 functions mostly -but not exclusively- in restricting/repressing gene expression either directly (Fig. 2D) or indirectly (Fig. S2C), likely as part of the gene regulatory networks (GRNs) that control early forelimb bud development (reviewed by Zuniga and Zeller, 2020).

### *Tbx3* regulates the spatial expression domains of diverse transcriptional regulators with key functions during early limb bud development

Next, the fraction of TBX3 candidate target genes with known spatial expression patterns and/or essential functions during limb development were annotated (Fig. 3A, 3B and Table S5). This analysis identified 19 genes whose expression is positively regulated (Fig. 3A), while 34 genes are inhibited/repressed by TBX3 in wildtype limb buds (Fig. 3B). Furthermore, this gene annotation analysis (Table S5) allowed construction of a TBX3-target GRNs for early limb buds (E10.0-E10.5) with several interesting features (Fig. 3C). This GRN reveals that the majority of annotated TBX3 target genes expressed in the anterior and proximal mesenchyme and AER are repressed by TBX3. This is also the case for several genes expressed in the distal mesenchyme, while most of the TBX3 target genes expressed in the posterior mesenchyme are positively regulated. Moreover, the majority of the annotated TBX3 target genes are TFs (30 of 43 genes; indicated by asterisks, Fig. 3C), which significantly enlarges the TF GRN and interactions that function during the onset of limb bud development (reviewed by Zuniga and Zeller, 2020).

**Figure 3.**
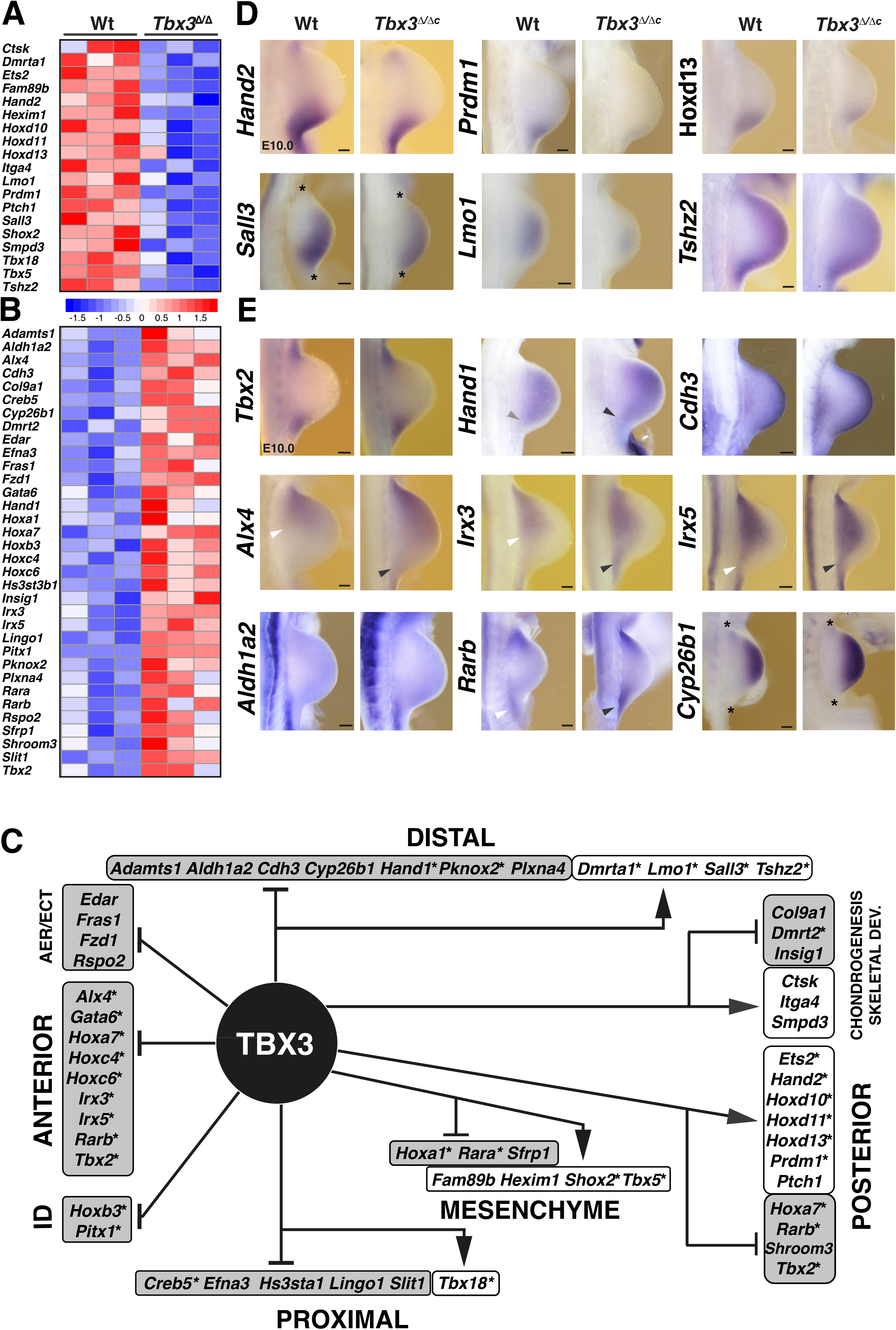
TBX3 candidate target gene regulatory network in early limb development. (A, B) Heatmaps illustrating relative gene expression of developmental regulator genes that are candidate TBX3 targets in wildtype and *Tbx3*-deficient samples (n=3 biological replicates, E9.75-10.0). The z-score scale represents mean-subtracted regularized log-transformed read counts. Shown are genes (n=53) that have essential functions in limb buds (Table S5). (C) TBX3 target gene regulatory network constructed by considering the limb bud expression patterns along the antero-posterior and proximo-distal limb bud axes; in the mesenchyme and AER/ectoderm (ECT). In addition, target genes with specific functions in limb identity (ID) or chondrogenesis and skeletal development are shown. Arrows point to groups of genes, whose expression is positively regulated by TBX3, while the inhibitory arrows indicate groups of genes whose expression is repressed by TBX3. Asterisks indicate transcription factors. (D-E) Comparative WISH analysis of select genes in wildtype and *Tbx3*^1/1c^ mouse forelimb buds at E9.75-10.25 (29-32 somites) of genes whose transcript levels are either down-regulated (panel D) or up-regulated in mutant forelimb buds (panel E). Asterisks indicate limb bud margins for some limb buds (panel D). Arrows heads indicate the posterior expression boundaries (*Alx4*; *Irx3*, *Irx5* or upregulated expression (*Hand1*) in wildtype (white) and mutant limb buds (black). All forelimb buds are oriented with anterior to the top and posterior to the bottom. Scale bars: 100 µm.

To assess spatial changes in target gene expression, *Tbx3* was inactivated in the mouse limb bud mesenchyme using the *Prxx1*-CRE driver (Logan et al., 2002) in mouse embryos carrying the *Tbx3*^ΔVenus^ null (*Tbx3*^Δ^) and a conditional *Tbx3*^flox^ allele (Fig. S3A, Frank et al., 2012). Whole mount immunofluorescence analysis of *Tbx3*^1/1c^ forelimb buds shows that the TBX3 protein is cleared from the anterior and posterior mesenchyme by E10.0, while it remains in the AER (Fig. S3B). Of note, fluorescent whole mount in situ by hybridization chain reaction (HCR^TM^) hybridisation detects variable levels of the non-functional *Tbx3*^ΔVenus^ transcripts in *Tbx3*^1/1c^ forelimb buds (E10.5, Fig. S3B). This conditional inactivation causes pre-axial duplications of digit 1 in combination with hypoplasia or loss of posterior digit 5. In addition variable defects affecting proximal limb skeletal elements are detected (ulna, radius and humerus, Fig. S3C, Emechebe et al., 2016).

Whole mount RNA *in situ* (WISH) analysis of specific mesenchymal TBX3 targets reveals that not only transcript levels (Fig. 3A, 3B) but also the spatial expression boundaries of several genes are altered in *Tbx3*^1/1c^ forelimb buds at E10.0 (Fig. 3D,E). For example, the reduced expression of several posterior target genes (Fig. 3A) is paralleled by more posteriorly (*Hand2*, *Prdm1*, *Hoxd11*, *Hoxd13, Ets2*) or distally (*Sall3* and *Lmo1*) restricted domains while *Tshz2* expression is more diffuse in *Tbx3*^1/1c^ than in wildtype forelimb buds (Fig 3D, Fig. 5C for *Ets2*). Next, we analysed the spatial expression of target genes that are up-regulated in *Tbx3* -deficient forelimb buds (Fig. 3B, 3E). Even though transcriptional up-regulation is more difficult to detect than down-regulation (Gamart et al., 2021), upregulation and spatial alterations are detected for several target genes in *Tbx3*^1/1c^ forelimb buds (Fig. 3E). In particular, the expression of several transcriptional regulators expressed highest in the anterior mesenchyme, namely *Hand1*, *Alx4*, *Irx3* and *Irx5*, is posteriorly expanded in mutant forelimb buds (arrowheads, Fig. 3E). Functional annotation of target genes shows that TBX3 negatively regulates key components of the retinoic acid (RA) pathway in early limb buds, namely the RA synthesising enzyme *Aldh1a2 (Raldh2)*, the receptors *Rara* and *Rarb* and the RA-degrading enzyme *Cyp26b1* which are all up-regulated in *Tbx3*^1/1c^ forelimb buds (Fig. 3B, 3C and Table S5). However, only two of these genes, *Rarb* and *Cyb261b* are expressed in the mesenchyme of wildtype forelimb buds while *Aldh2* is expressed in the posterior flank mesenchyme at E10.0 (lower panel, Fig. 3E). *Rarb* is upregulated and its expression boundary changed in the posterior-proximal mesenchyme (arrowheads, Fig. 3E). Concurrently, the *Cyp26b1* expression is less distally restricted, which indicates that RA pathway activity could be altered in *Tbx3*^1/1c^ forelimb buds. Taken together, these results show that TBX3 preferentially downregulates/represses target genes in the anterior and distal mesenchyme, while most posterior target genes are positively regulated (Fig. 3C). The posterior expansion of *Hand1, Alx4, Irx3/5* and enlargement of *Rarb* domain (arrowheads, Fig. 3E) points to a specific TBX3 repressor function in posterior mesenchymal expression boundary formation.

### Shared target genes reveal distinct roles for *Tbx3* and *Hand2* in regulating posterior identity

Previous genetic and molecular analysis showed that *Tbx3* and *Hand2* participate in restricting *Gli3* expression from the posterior mesenchyme, but did not resolve the underlying genetic and molecular hierarchies and extent to which these two TFs impact the spatial expression of other target genes during the onset of forelimb bud development. As *Tbx3* and *Hand2* are both required to up-regulate each-other’s expression in the posterior limb bud mesenchyme (Fig. 3A, 3D), this is indicative of feed-forward regulation. To gain insight into the functional importance of the *Tbx3*-*Hand2* interactions, the skeletal phenotypes of mouse embryos lacking *Hand2*^1/1c^,*Tbx3*^1/1c^ and *Tbx3* ^1/1c^, *Hand2* ^1/+^ in forelimb buds were comparatively analysed at E14.5 (Fig. 4A-C). *Hand2*^1/1c^ forelimb buds (Fig. 4A) display the characteristic distal limb skeletal truncations and absence of the ulna (Fig. 4A), and *Tbx3* expression is absent from the posterior mesenchyme of *Hand2-*deficient limb buds by E10.0 (Fig. 4D, Osterwalder et al. 2014). In contrast, *Tbx3*^1/1c^ forelimbs show duplications of distal phalanges and associated metacarpals of the anterior digit 1 in combination with digit 5 hypoplasia or agenesis (digit phenotypes indicated by asterisks, Fig. 4B, Emechebe et al., 2016). In addition, variable degrees of agenesis of the deltoid tuberosity, ulnar hypoplasia and thickening of the radius are detected (Fig.4B, Lopatka and Moon, 2022). *Hand2* remains expressed in the posterior mesenchyme of *Tbx3*^1/1c^ forelimb buds, but levels are reduced (Fig. 4D, 4F). Genetic inactivation of one *Hand2* allele in *Tbx3*-deficient limbs results in more severe skeletal defects that include loss of metacarpals and digits (Fig. 4C). The most severe phenotypes bear striking similarities to the *Hand2* loss-of function phenotypes with exception of maintaining the digit1 duplications, which are a hallmark of *Tbx3*-deficient forelimbs (lowest panel Fig. 4C, compare to Fig. 4B). This genetic interaction analysis shows that *Hand2* and *Tbx3* interact in the posterior limb bud mesenchyme, while the *Tbx3* specific functions in the anterior mesenchyme are not linked to *Hand2* (Fig. 4D), which is consistent with its spatial expression and essential role in regulating GLI3 protein stability (Emechebe et al., 2016). As *Tbx3* expression is lost from the posterior mesenchyme of *Hand2*-deficient limb buds, the comparative analysis of *Tbx3*^1/1c^ and *Hand2*^1/1c^ forelimb buds should identify the *Tbx3*-specific and shared *Tbx3*-*Hand2* functions in target gene regulation.

**Figure 4.**
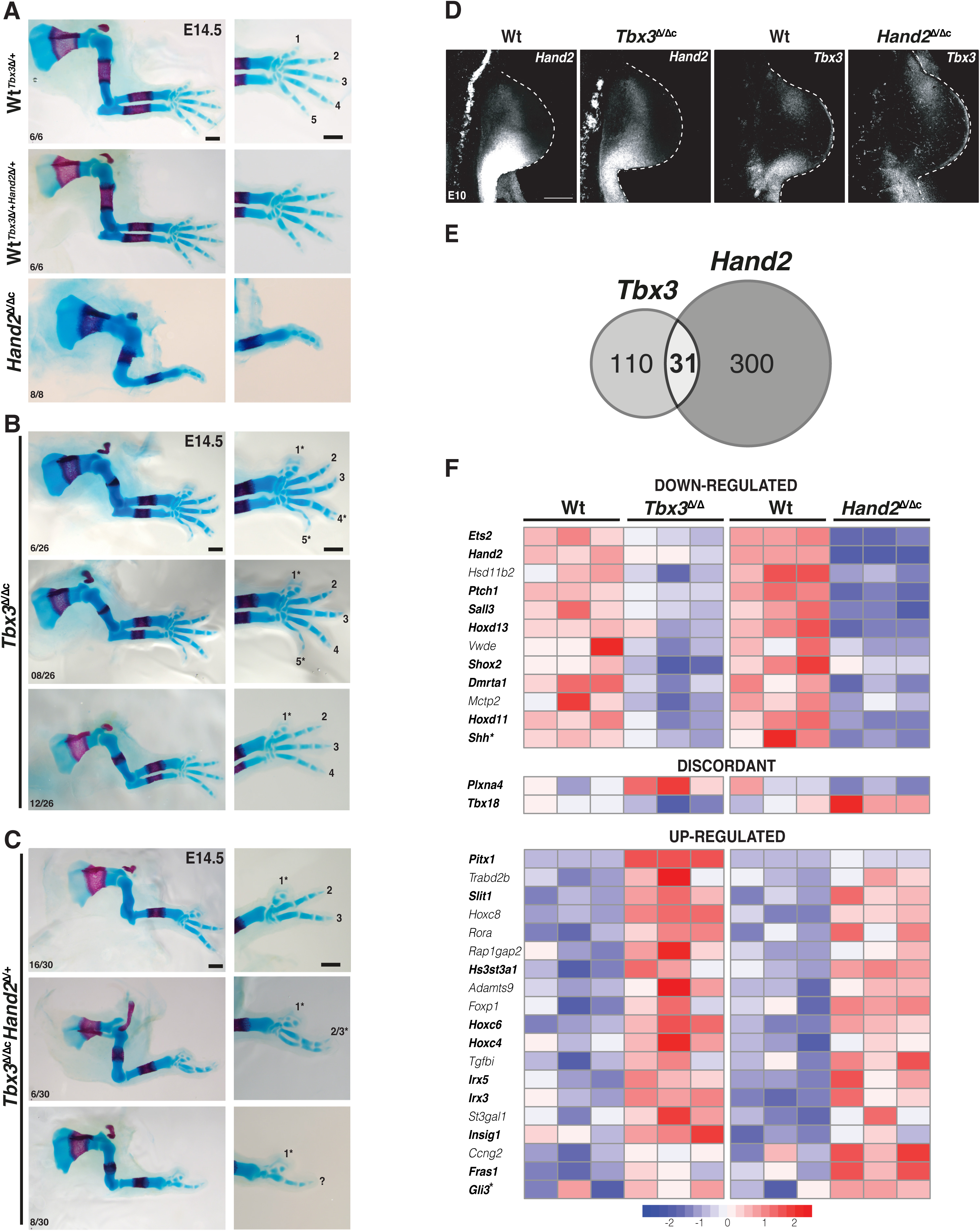
*Tbx3* and *Hand2* cooperatively control posterior limb skeletal identities and co-regulate shared target genes in early limb buds. (A-C) Comparative skeletal analysis of wildtype, *Hand2*-deficient (panel A), *Tbx3*^1/1c^ (panel B) and *Tbx3*^1/1c^Hand2^1/+^ (panel C) mouse forelimbs at E14.5. This analysis reveals the genetic interaction of *Tbx3* and *Hand2* in regulation of posterior limb skeletal elements (ulna and posterior digits, panel C). Scale bars: 500µm. Digit identities based on digit position and shape are indicated from anterior to posterior (1-5). (D) HCR^TM^ analysis of forelimb buds at E10.0 (29-32 somites). *Hand2* expression is reduced in *Tbx3*^1/1c^ forelimb buds, while the posterior *Tbx3* expression is lost *Hand2*^1/1c^ forelimb buds (n≥4 biological replicates per genotype). All limb buds are oriented with anterior to the top and posterior to the bottom. Scale bar: 200µm. (E) TBX3 and HAND2 share a small subset of their target genes (n=31). (F) Heatmap of the common target genes that are co-regulated by TBX3 and HAND2. Genes in bold are genes with known functions during early limb bud development. Asterisks indicate the fact that *Shh* and *Gli3* were manually curated and added to the list of target genes.

The HAND2 target genes in early limb bud were identified by reanalysing the ChIP-seq dataset by Osterwalder et al. (2014) in combination with the differentially expressed genes in wildtype and *Hand2*-deficient forelimb buds (E10.0, n=3 biological replicates per genotype, Table S7). This analysis identified 331 differentially expressed HAND2 candidate target genes with a fold change ≥1.2 (p-value ≤0.05, Fig. S4 and Table S8). In *Hand2*-deficient forelimb buds 124 target genes are down- and 209 up-regulated (Fig. S4). The common TBX3 and HAND2 target genes were identified by overlapping the *Tbx3* and Hand2 datasets, which identified 31 shared target genes that are DEGs in both *Tbx3* and *Hand2* mutant limb buds (E10.0, Fig. 4E, 4F and Table S9). Two additional genes, *Shh* and *Gli3* known to critically depend on TBX3 and HAND2 were added to the list of target genes by manual curation (asterisks in Fig. 4F and Table S9) for the following reasons: *Shh* has been previously identified as a HAND2 target gene (Galli et al., 2010) and its expression is significantly down-regulated in both *Tbx3* and *Hand2* mutant limb buds (Table S9). The association of the TBX3 ChIP-seq peaks located in the distant *Shh* limb bud enhancer called ZRS (Fig. S5) with *Shh* failed due to the large distance between this upstream enhancer and the *Shh* promoter (≥800Mb, Lettice and Hill, 2005). *Gli3* is a HAND2 target gene (Osterwalder et al., 2014) and in this study we establish that TBX3 ChIP-seq peaks are detected in several *Gli3* enhancers (see below). Nevertheless, *Gli3* was not scored as DEG by RNA-seq of early forelimb limb buds as the *Gli3* upregulation falls just below the threshold (>1.2), with a 1.15-fold upregulation in*Tbx3*^1/1c^ and 1.2-fold increase in *Hand2*^1/1c^ forelimb buds (Table S9).

Of all candidate target genes, just two are regulated in a discordant manner by TBX3 and HAND2 (middle panel, Fig. 4F). All other targets are concordantly controlled by both TFs with their expression being either activated/enhanced or repressed/reduced in early forelimb buds. Nine of the 12 target genes downregulated in both *Tbx3*^1/1c^ and *Hand2*^1/1c^ forelimb buds are expressed in the posterior and/or distal mesenchyme (in bold, top panel of Fig. 4F). These positively regulated genes function either upstream or downstream of the SHH pathway in early limb buds. Furthermore, 10 of 19 target genes up-regulated in both mutants have distinct spatial expression patterns and/or essential functions in early limb buds (in bold, bottom panel of Fig. 4F). Eight of these are expressed in the proximal (*Hs3st3a1*, *Pitx1*, *Slit1*) or anterior/anterior-distal limb bud mesenchyme (*Hoxc4*, *Hoxc6*, *Irx3*, *Irx5*, *Gli3*). Taken together, this analysis shows that *Hand2* and *Tbx3* coregulate major TFs in the GRN that orchestrates limb axes patterning and restricts establishment of the posterior *Shh* signaling centre.

Next, the specific and shared effects of HAND2 and TBX3 on spatial expression of target genes was assessed by fluorescent HCR^TM^ whole mount *in situ* hybridisation, which allows for precise co-localisation of gene expression. TBX3 and HAND2 directly impact the spatial expression of *Shh* and its receptor *Ptch1* (Fig. 5A). In *Tbx3*^1/1c^ forelimb buds, both *Shh* and *Ptch1* are downregulated and their domains are more posteriorly restricted (top and middle panels, Fig. 5A). In *Hand2*^1/1c^ forelimb buds much lower *Shh* and *Ptch1* expression is detected (lower panels, Fig. 5A, Galli et al., 2010). As HAND2 and TBX3 are required for activation and upregulation of *Shh* expression, respectively, the concurrent loss of *Ptch1* is foremost indicative of disrupting SHH signalling in *Hand2*^1/1c^ forelimb buds. In addition to HAND2, posterior *5’Hox* genes including *Hoxd11* and *Hoxd13* function in *Shh* activation (Kmita et al., 2005), while *Ets2* functions downstream of *Hand2* and *Tbx3* to upregulate *Shh* expression (Lettice et al., 2012). In *Tbx3*^1/1c^ forelimb buds, the spatial domains of *Hoxd11/13* and *Ets2* are slightly more restricted (top and middle panels Fig. 5B, 5C) while they are restricted to a small posterior domain in *Hand2*^1/1c^ forelimb buds (lower panels, Fig. 5B, 5C). The drastic reduction of posterior *Hoxd* gene expression points to feedback regulation between these genes and *Hand2*. Together these data indicate that HAND2 functions in activation/initial upregulation of posterior target genes, while TBX3 functions in enhancing gene expression.

**Figure 5.**
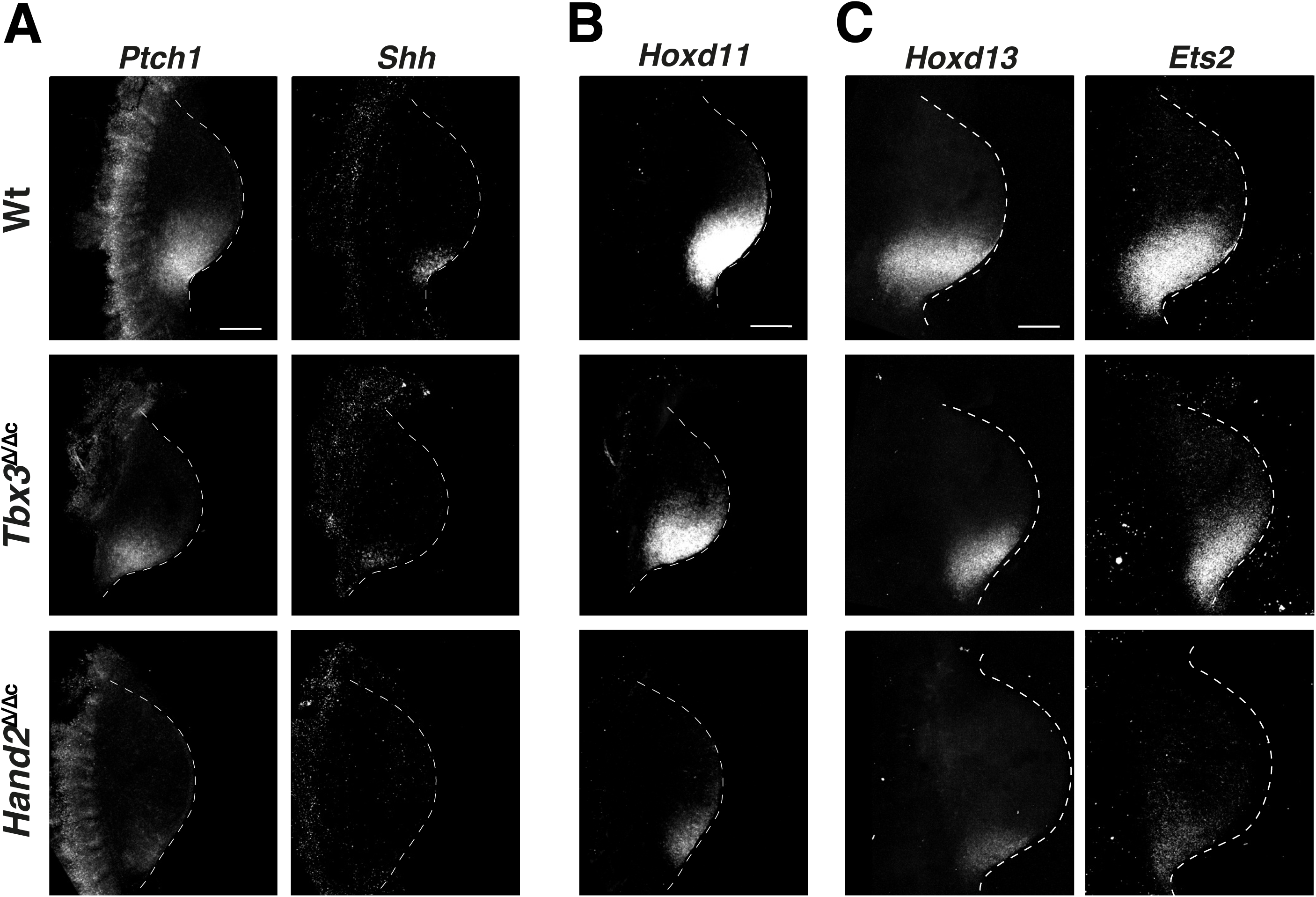
*Hand2* and *Tbx3* requirement for activation and spatial up-regulation of key regulators functioning in establishment of posterior identities. (A) Comparative HCR^TM^ analysis of the spatial alterations in the expression of the shared target genes *Ptch1* and *Shh* in wildtype, *Tbx3*^1/1c^ and *Hand2*^1/1c^ forelimb buds at E10.0 (29-32 somites). For all three genotypes the two genes were co-localized using different fluorophores in the same forelimb bud. (B,C) Comparative HCR^TM^ analysis of the transcriptional regulators *Hoxd11* (panel B), *Hoxd13* and *Ets2* (panel C) in wildtype and mutant forelimb buds. *Hoxd13* and *Ets2* were co-localised in the same limb bud, while the *Hoxd11* analysis was done in different limb buds. All forelimb buds are oriented with anterior to the top and posterior to the bottom. For all genes n≥4 biological replicates per genotype were analysed. Scale bar: 200µm.

Next, potential changes in the spatial expression of shared target transcript factors expressed in the distal and anterior/proximal limb bud mesenchyme were assessed. In particular, HCR^TM^ whole mount *in situ* hybridisation showed that posterior *Tbx3* and anterior-proximal *Irx3* are expressed in complementary patterns in early forelimb buds, while the *Tbx3* and anterior-distal *Sall3* expression domains show a small overlap in their posterior boundary region (E10.0, top panels, Fig. 6A). In *Tbx3*^1/1c^ forelimb buds, the spatial domain of *Irx3* expands to the posterior limb bud margin (*Irx3*, middle panel, Fig. 6A), which contrasts with the lack of posterior expansion of the *Sall3* domain (*Sall3* middle panel, Fig. 6A). This is interesting in light of the fact that there is no distinct posterior boundary between the *Tbx3* and *Sall3* expression domains in wildtype forelimb buds (top panels, Fig. 6A). In addition, the proximo-distal boundaries between the *Irx3* and *Sall3* expression domains is maintained (middle panel, Fig. 6A). In *Hand2*^1/1c^ forelimb buds that lack posterior *Tbx3* expression (Fig. 4D), no additional alterations in the spatial expression of *Irx3* and *Sall3* are detected (bottom panel, compare to middle panel, Fig. 6A). These results indicate that the posterior expression boundary of *Irx3* (but not *Sall3*) is regulated by TBX3-mediated transcriptional repression. To substantiate this assumption, we analysed the anterior-proximal expression domain of *Irx5* and the anterior-distal expression domain of *Gli3* (Fig. 6B) which are upregulated in mutant forelimb buds (Fig. 4F). Their spatial expression boundaries are complementary and nonoverlapping with *Tbx3* in the posterior mesenchyme (top panels, Fig. 6B). Both *Irx5* and *Gli3* expression expands to the posterior margin in *Tbx3*^1/1c^ and *Hand2*^1/1c^ forelimb buds, while their proximo-distal complementarity is maintained (middle and bottom panels, Fig. 6B and Fig. S6 for *Gli3*). These results uncover a specific requirement for TBX3 in posterior gene expression boundary formation of anterior genes during AP limb axis polarisation. The results shown in Figs. 5 and 6 establish that TBX3-dependent boundary formation excludes both anterior-proximal and anterior-distal genes from the posterior mesenchyme in which the SHH signalling limb organiser is subsequently established. Together this analysis reveals (a) the critical repressive function of TBX3 in establishing the posterior expression boundaries of several anteriorly expressed transcriptional regulators (Fig. 4E and Fig. 6) and that (b) TBX3 participates in transcriptional upregulation of posterior identity genes together with HAND2 (Fig. 5).

**Figure 6.**
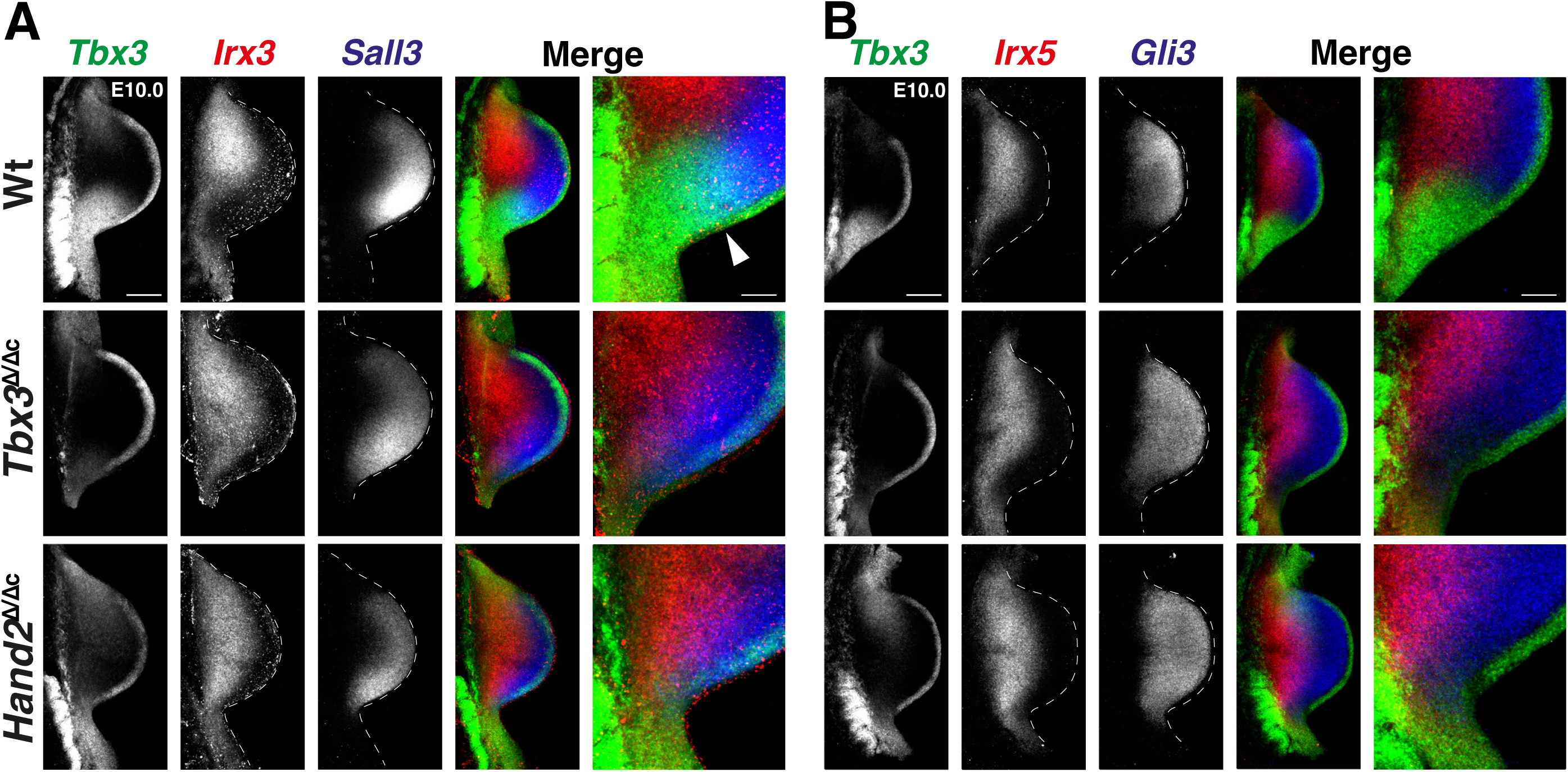
An unexpected essential TBX3 function in posterior expression boundary formation is independent of HAND2. (A) Co-localization of the spatial *Tbx3* (green)*, Irx3* (red)*, Sall3* (blue) expression in forelimb buds (E10.0,29-32 somites) of all three genotypes (wildtype, *Tbx3*^1/1c^ and *Hand2*^1/1c^). Minimally n=4 biological replicates were analysed per gene and genotype. The right-most panels show enlargements of the posterior proximal regions. The white arrowhead points to the region of overlap between *Tbx3* (green) and *Sall3* (blue) expression, while no such overlap is seen between *Tbx3* and *Irx3*, and *Irx3* and *Sall3* in wildtype (Wt) forelimb buds. (B) Co-localization of the spatial *Tbx3* (green)*, Irx5* (red) and *Gli3* (blue) expression in forelimb buds at E10.0. Minimally n=3 biological replicates were analysed per gene and genotype. In the enlargements of the posterior proximal regions, clear boundaries with no overlaps demarcate the spatial domains of all three genes in wildtype limb buds. Note the specific loss of the posterior *Tbx3* expression in *Hand2*^1/1c^ forelimb buds. All limb buds are oriented with anterior to the top and posterior to the bottom. Scale bar: 200µm and 67 µm for the enlargements.

### TBX3 controls posterior gene expression boundary formation by direct repression of enhancer activities

The anterior-posteriorly graded *Gli3* expression in mouse limb buds (Fig. 6B) is controlled by two functionally redundant enhancers embedded in the large *Gli3* genomic landscape (Osterwalder et al., 2018). This provides a unique opportunity to determine the extent to which TBX3 repression might directly impact the spatial activities of these two enhancers. Initially, the spatial activities of three *Gli3* CRMs encoding genomic regions enriched in both TBX3 and HAND2 (VISTA mouse enhancer *mm1179*) or specifically TBX3 chromatin complexes (VISTA mouse orthologue enhancer *mm-hs1586* and *mm652*) were assessed by mouse embryonic *LacZ* reporter assays (Fig. S7, Visel et al., 2007). Activity of the *mm652* enhancer is detected transiently in a fraction of transgenic founder embryos during early hindlimb bud development (E10.5, right-most panel, Fig. S7A) and was not analysed further. In contrast, the functionally redundant *mm1179* and *mm-hs1586* enhancers possess robust activity in forelimb buds (upper panels, Fig. S7B, Osterwalder et al., 2018). Therefore, the most conserved bases in the predicted TBX3 and HAND2 binding sites were mutated to disrupt TF binding (Fig. S7C and Fig. S8). *LacZ* reporter analysis shows that the activity of both mutant enhancers is more restricted but fails to expand posteriorly (Fig. S7B). This shows that the activity of these two *Gli3* enhancers in mouse limb buds likely depends on a combination of both positive and repressive trans-regulatory inputs from different T-box family transcription factors. The repressive effects on the *hs-mm1586* limb bud enhancer is apparent from ectopic activity of the mutant enhancer in eye primordia and facial tissues co-expressing *Tbx3* and *Gli3* (Fig. S7D).

As this approach was not informative with respect to the specific functions of TBX3 in posterior boundary formation in limb buds, we generated stable transgenic reporter mouse strains for the *mm1179* and *mm-hs1586* enhancers (Fig. 7A) that reproduce their spatial activities. These *LacZ* reporter transgenes were then crossed into *Tbx3*^1/1c^ limb buds and analysed in comparison to wildtype controls by HCR^TM^ whole mount *in situ* hybridisation (Fig. 7B-K). The activities of both enhancers are restricted to the posterior boundary set by the *Tbx3* expression domain in wildtype limb buds, but Their activities expand to the posterior margin in *Tbx3*-deficient forelimb buds (*mm1179*: compare Fig. 7B, 7C, 7E; *mm-hs1586*: Fig. 7G, 7H, 7J and online movies 1, 2). The posterior expansion of enhancer activity perfectly matches the expansion of *Gli3* expression in *Tbx3*^1/1c^ forelimb buds (*mm1179*: compare Fig. 7C, 7D, 7F; *mm-hs1586*: Fig. 7H, 7I, 7K). This together with the TBX3 ChIP-seq peaks in the core regions of both enhancers (Fig. 7A) indicates that TBX3 binds these enhancers to repress *Gli3* in the posterior mesenchyme. In *Tbx3*-deficient forelimb buds, the enhancer activity is not only expanded but also increase in comparison to wildtype limb buds (Fig. 7C, 7H), which underscores the strong repressive effect of TBX3. In summary, this analysis reveals that TBX3 controls posterior expression boundary formation of *Gli3* and several additional TFs such as *Alx4*, *Hand1*, *Rarb* and *Irx3/5* by repressing their expression from the posterior-proximal limb bud mesenchyme (Fig. 3E, Fig. 6 and Fig. S6).

**Figure 7.**
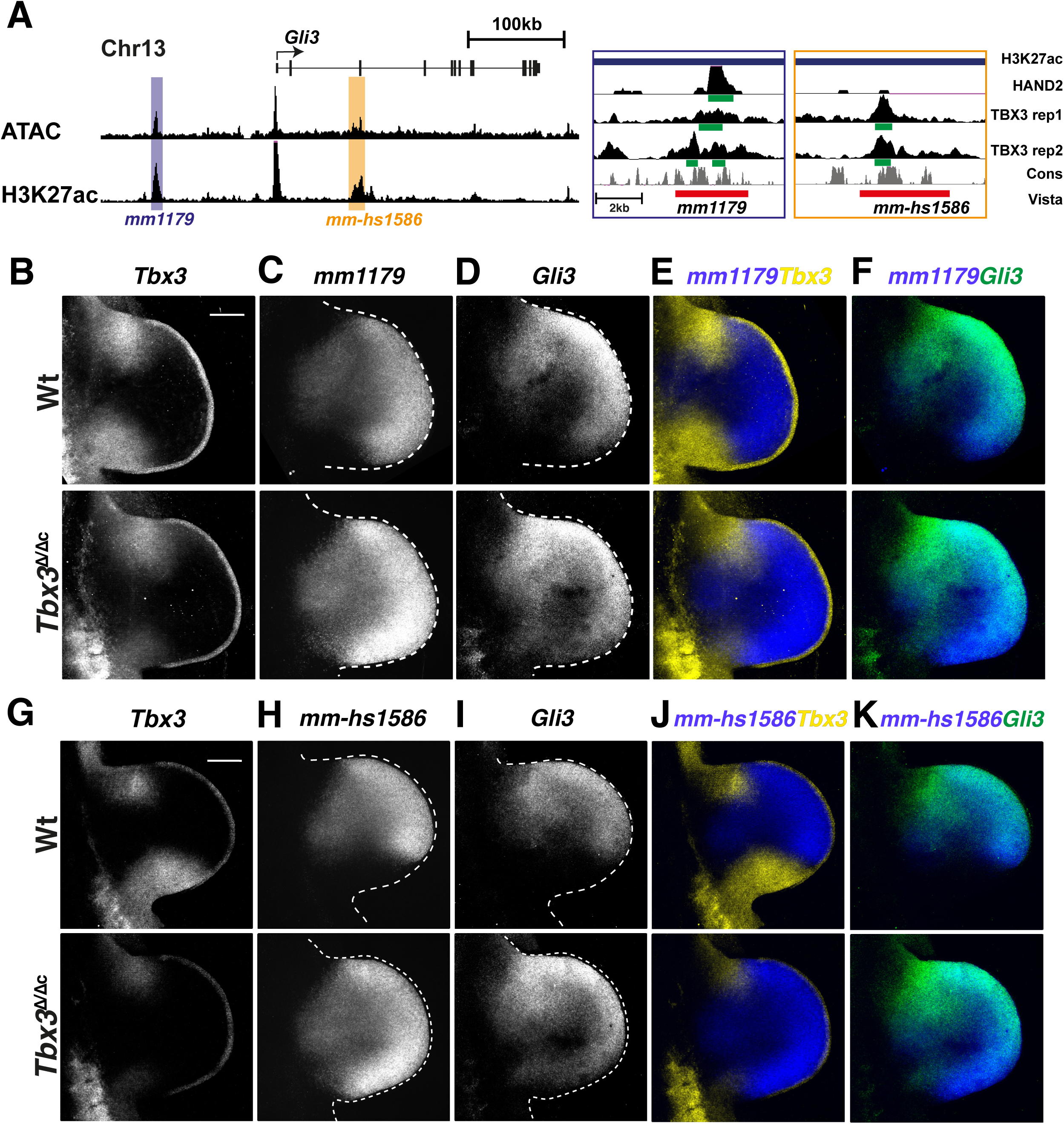
*Tbx3* is required to restrict the activity of the *Gli3* limb enhancers and *Gli3* expression from the posterior limb bud mesenchyme. (A) UCSC browser view of parts of the *Gli3* topologically associating domain with regions of accessible and active chromatin (ATAC-seq and H3K27ac peaks). The two limb enhancers mm1179 and mm-hs1586 are highlighted. The enlargements of the genomic regions for these two enhancers (right panel) show the genomic region required for *Gli3* enhancer activity (red bar) and the HAND2 and TBX3 ChIP-seq peaks mapping to these enhancers. The green bars indicate the called peaks. Note that both replicates for the TBX3 ChIP-seq are shown, while the HAND2 ChIP-seq data is from Osterwalder et al., 2014. (B-K) Comparative HCR^TM^ analysis of wildtype (upper panels) and *Tbx3*^1/1c^ (lower panels) mouse limb buds at E10.5 (35–37 somites) shows the spatial expression of endogenous *Tbx3* (yellow), transgenic enhancer-*LacZ* reporter RNA expression (blue; mm1179: panels C,E,F and mm-Hs1586: panels H, J, K) and endogenous *Gli3* expression (green). Minimally n=5 biological replicates were analysed for probes and genotypes. Note: low and variable levels of the protein-null *Tbx3* ^Δ^ transcript can be detected in *Tbx3*^1/1c^ limb buds due to the HCR^TM^ probe set. This is apparent in the *Tbx3*^1/1c^ forelimb buds shown in panels B, E (anterior and low posterior) and G, J (anterior only). All forelimb buds are oriented with anterior to the top and posterior to the bottom. Scale bars: 200µm.

## DISCUSSION

The transcriptional repression of *Gli3* hallmarks the establishment of the posterior mesenchymal territory competent to activate *Shh* expression (Litingtung et al., 2002; Welscher et al., 2002b). This requires the interaction of HAND2 and several homeodomain transcriptional regulators (5’HOXD and PBX) with the distant ZRS *Shh* enhancer (Capellini et al., 2006; Galli et al., 2010; Losa et al., 2023). To gain insight into the underlying gene regulatory logic in TBX3 functions during establishment of AP polarity, we identified its direct transcriptional targets in the mesenchyme of early forelimb buds. The fraction of target genes upregulated in *Tbx3*-deficient limb buds (75%) reveals the predominant role of TBX3 as a transcriptional repressor. It is known that TBX3 directly binds to *T-box* motifs and represses transcription by interaction of its carboxy-terminal repressor domain with HDAC histone deacetylases (Yarosh et al., 2008). During early forelimb bud development, *Tbx3* is first expressed posteriorly as its activation in the anterior mesenchyme is temporally delayed (Emechebe et al., 2016, this study). The TBX3 target GRN reveals two distinct features of gene regulation at early stages: (1) the expression of several anteriorly expressed TBX3 target genes is repressed from the posterior limb bud mesenchyme; (2) TBX3 target genes whose expression is positively regulated include a large fraction of TFs functioning in establishment of posterior identity and *Shh* expression. In particular, *Tbx3* is required to up-regulate *5’Hoxd* genes, *Hand2* and *Ets2* that in turn function to activate and/or upregulate *Shh* expression in the posterior mesenchyme (Kmita et al., 2005; Galli et al., 2010; Lettice et al., 2012). As TBX3 acts predominantly as a repressor (Carlson et al., 2001) positive regulation of target genes must depend on interactions with other transcriptional regulators. A recent study established that TBX3 interacts with BCL9 as part of WNT/ß-Catenin transcriptional complexes, which up-regulates gene expression during forelimb bud outgrowth (E10.5, Zimmerli et al., 2020). Indeed, WNT signaling has been implicated in formation of chicken limb buds (Kawakami et al., 2001), but to date there is no genetic evidence for an equivalent role in early mouse limb buds. It is also possible that TBX3 interacts with other TBX transcriptional regulators as is the case during mouse kidney development, where TBX2 and TBX3 function downstream of canonical WNT signal transduction during ureteric mesenchyme patterning and differentiation (Aydogdu et al., 2018). This could also be relevant to limb bud development, as *Tbx2* and *Tbx3* are co-expressed in the anterior and posterior mesenchyme and the spatial expression of *Tbx2* is not altered in *Tbx3* mutant limb buds (this study). Indeed, the stepwise reduction of the *Tbx2* and *Tbx3* gene dosage increases the severity of the limb skeletal malformations, which are not strictly additive but in parts synergistic (Lopatka and Moon, 2022). As the core sequence of the T-box motif enriched in the TBX3 cistrome is shared by different TBX proteins, coregulation of target genes by several *Tbx* genes expressed in limb buds is likely. Spatial regulation of target genes may also involve *Tbx18, Tbx15* and *Tbx5*, which are expressed in a rather complementary fashion to *Tbx2/*3 in early limb buds (Sheeba and Logan, 2017). In particular, their expression patterns significantly overlap the spatial activity patterns of the *Gli3* enhancers and *Gli3* transcripts. Finally, *de novo* motif analysis reveals enrichment of other types of TF motifs in the TBX3 cistrome, which may participate with *Tbx3* in spatial regulation. For example, HOX and TBX TFs extensively co-bind *T-box*-*Hox* composite motifs in the mouse genome during limb bud development, which serves to integrate different inputs into target gene regulation (Jain et al., 2018).

Most importantly, this study establishes that *Tbx3* functions downstream of *Hand2* and is the transcription factor required to establish the posterior expression boundaries of anterior-proximal and anterior-distally expressed genes. In wildtype limb buds, sharp posterior and non-overlapping expression boundaries with *Tbx3* are observed for *Gli3*, *Alx4*, *Hand1*, *Irx3* and *Irx5*, whilst their expression expands to the posterior flank in both *Tbx3*^1/1c^ and *Hand2*^1/1c^ limb buds. Comparative analysis of *Gli3* enhancer activities and *Gli3* expression reveals the precision by which TBX3 controls posterior expression boundary formation via the two enhancers that provide *Gli3* expression with robustness(this study and Osterwalder et al., 2018). The results of the present study are consistent with previous analysis of other loss-of function mutations that act upstream of *Hand2* and *Tbx3.* Inactivation of all *Hox9* paralogues and *Isl1* disrupts *Hand2* expression in fore- and hindlimb buds, respectively, which results in posterior expansion of *Gli3* expression and lack of *Shh* expression (Xu and Wellik, 2011; Itou et al., 2012 and this study). The TALE homeodomain TFs *Meis1* and *Meis2* are required for proximo-distal limb bud axis patterning, but genetic analysis reveals an additional early requirement of *Meis1/2* in AP limb bud axis patterning (Delgado et al., 2020; Delgado et al., 2021). Both MEIS TFs are required for upregulation of *5’Hoxd* expression and activation of *Hand2* expression in the posterior limb bud mesenchyme. In particular, MEIS proteins interact with an enhancer that is required for *Hand2* expression (Delgado et al., 2021). This *Hand2* enhancer is also enriched PBX chromatin complexes in agreement with *Pbx1/2* requirement for *Hand2* upregulation in mouse limb buds (Capellini et al., 2006; Losa et al., 2023). In forelimb buds lacking both *HoxA* and *HoxD* gene functions, *Hand2* remains expressed while *Shh* activation is disrupted (Kmita et al., 2005), which indicates that *HoxA* and *HoxD* genes function downstream of *Hand2 and Tbx3* (this study). These studies reveal the genetic hierarchies and molecular interactions that converge on activation and upregulation of *Hand2* expression, which is in turn required to activate *Tbx3* expression in the posterior limb bud mesenchyme. As the *Hand2* expression domain is larger than the one of *Tbx3*, other transcriptional regulators must contribute to define the spatial *Tbx3* expression domain in the posterior mesenchyme. The importance of generating these expression boundaries and the posterior mesenchymal territory has been corroborated by several studies showing that initially *Hand2* (via *Tbx3*) and subsequently SHH (by inhibition of GLI3 processing) overcome the “ground state” GLI3R repression in the early limb mesenchyme to enable posterior identities and *Shh* activation (Welscher et al., 2002b; Osterwalder et al., 2014). The establishment of this posterior GLI3R-free territory enables activation of SHH signaling, which patterns the future digits at an early stage and promotes distal limb bud outgrowth (Litingtung et al., 2002; Welscher et al., 2002b; Zhu et al., 2008; Zhu et al., 2022).

The present study unveils a previously unrecognized essential role of TBX3 in shaping the posterior expression boundary of anterior genes including *Irx3/5*. Interestingly, the *Drosophila Tbx2/3* orthologue *Optomotor blind* (*Omb*) is expressed in a complementary manner to the *Drosophila Irx* orthologue *Iroquois-C* (*Iro-C*) in the wing imaginal disk, which separates the wing hinge and notum territories (Wang et al., 2016). Loss- and gain-of-function analysis by Wang and co-workers (2016) showed that *Omb* directly represses *Iro-C*, which is both necessary and sufficient for formation of the boundary fold between the hinge and notum in the wing imaginal disc. Thus, the repression of *Irx3* and *Irx5* by TBX3 is an evolutionary ancient gene regulatory circuit required for insect wing and leg imaginal disc development (Pflugfelder et al., 2017) that has been co-opted to the establishment of AP axis polarity and the posterior mesenchymal territory in vertebrate limb buds.

## MATERIALS AND METHODS

### Ethics statement, mouse strains and embryos

All experiments were conducted with the different *Tbx3* and *Hand2* alleles bred into a Swiss Albino (*Mus musculus*) as only robust phenotypes manifest in this strain background with large litter sizes (15-20 embryos), which is in line with the refine and reduce 3R principles. Embryos of both sexes at the developmental ages indicated were used for experimental analysis in accordance Swiss laws and the 3R principles. Embryos were age-matched by counting somites and comparing limb shape and size. All animal studies were evaluated and approved by the Regional Commission on Animal Experimentation and the Cantonal Veterinary Office of Basel (national license 1951). In addition to the *Tbx3*^ΔVenus^ null allele (Kunasegaran et al., 2014), the *Prrx1-Cre* strain (Logan et al., 2002) was used to conditionally inactivate the floxed *Tbx3* (Frank et al., 2013) and *Hand2* (Galli et al., 2010) alleles in the mouse forelimb bud mesenchyme. Wildtype and littermates carrying the *Prrx1-Cre* transgene were used as controls. Primers are listed in Table S10.

### Generation of the *Tbx3*^3xF^ mouse allele

The 3XFLAG epitope tag was inserted in frame into the carboxy-terminal exon of the *Tbx3* open reading frame. CRISPR/Cas9-mediated homology-directed repair in combination with a 200bp single-stranded DNA oligonucleotide (ssODNs) repair template that also encodes the sequence for the triple FLAG peptide. Mouse G4 ES cells were transfected with a mix containing the targeting vector that also encodes the Cas9 nuclease and its single guide (sg)RNAs together with ssODNs repair template. After recovery from transfection, cells were selected with 2mg/ml puromycin for 48hrs to enrich ES cells that have up-taken up DNA mix including the targeting vector. After picking and expanding 100-150 ES cell clones, correct genome editing events were detected by extensive PCR analysis. Correctly genome edited ES cell clones were further verified by sequencing the genomic region of the *Tbx3* locus carrying the 3xFLAG epitope insertion. Verified ES cell clones were used to generate several aggregation chimeras. Highly chimeric mice (≥70%) were then bread to germline and the genome-edited *Tbx3* verified again. Primers are listed in Table S10.

### Skeletal analysis

Embryos were collected at E14.5 into ice-cold PBS and fixed for 3-5 days in 95% ethanol. The ethanol was removed and the embryos stained for 24h with filtered Alcian Blue staining solution prepared by dissolving 30mg of Alcian Blue 8GX (Sigma-Aldrich, A3157) in 80ml of 95 % ethanol and 20ml of glacial acetic acid. Then they were rinsed twice for 15 min in 95% ethanol and stored for 24hrs in 95% ethanol. Then the samples are cleared in 1% KOH w/v in water for 90min to 3hrs. This is followed by counter-staining bone with Alizarin Red (50mg of Alizarin Red per liter 1% KOH) for 3hrs. Clearing is continued in 1% KOH solution for 30min, followed by changing the solution as follows to ratios v/v of 1%KOH/glycerol: 80/20 for 4 days or until the embryos have cleared, then the solution is changed every 48 hrs to 60/40, 40/60 and samples are stored tin 20/80 v/v 1%KOH/glycerol. Images were taken using a Leica MZ-FL2 stereo microscope. Per genotype ≥3 biological replicates were analyzed.

### Western blot experiments

Protein extracts were prepared from forelimb buds (E10.5, 33-36 somites) and 15mg of total proteins per sample loaded onto 12.5% SDS-PAGE gels. Following gel-electrophoresis proteins were transferred to a PVDF membrane (Merck, IPVH00010) using a semi-dry transfer cell (Biorad Transblot SD). TBX3^3xFLAG^ proteins were detected by chemo-luminescence using monoclonal M2 anti-FLAG antibodies (1:500, Sigma, F1804) in combination with donkey anti-mouse IgG-HRP antibodies (1:5000, AP192P, Millipore) using a ChemiDocTM XRS+ with Image Lab from Bio-Rad. TBX3 proteins were detected using rabbit anti-TBX3 antibodies (Abcam, 1:500, ab99302) with secondary goat anti-rabbit HRP antibodies (Millipore, 1:5000, AP187P).

### TBX3^3xF^ ChIP-seq analysis

Two independent biological replicates were analysed to ensure reproducibility following [ENCODE guidelines](https://www.encodeproject.org/about/experiment-guidelines/). For each replicate, forelimb buds of ∼70 *Tbx3^3xF/3xF^* mouse embryos at embryonic day E9.75-E10.25 (29 to 32 somites were dissected in ice-cold PBS and crosslinked in 1% formaldehyde for 10 min. Fixation of limb buds was stopped with glycine, limb buds washed in PBS (with protease inhibitors) and stored at -80°C. Frozen tissues were thawed on ice, nuclear extracts prepared and chromatin sheared by sonication to a size range of ∼200-500 base pairs. Per replicate 5mg of mouse-M2-anti-FLAG antibodies (F1804, Sigma) coupled to protein A/G beads were used for chromatin immunoprecipitation. After overnight incubation, the immunoprecipitated chromatin complexes were extensively washed, and the chromatin complexes eluted from the beads and the DNA purified after decrosslinking. After assessing the quality of the eluted DNA, libraries for sequencing were prepared using the MicroPlex library preparation kit v2 (C05010014, Diagenode). Libraries were purified using AMPure XP purification beads (A63880, Beckman Coulter Life Sciences) and sequenced using the Illumina NextSeq 500 system. A detailed ChIP-seq protocol is available on request.

### ChIP-seq analysis for identification of enriched regions

Paired-end reads of 75bp from TBX3 ChIP-seq experiments were subjected to quality check using FastQC v0.11.4 (Andrews, 2010). Trim_Galore v0.4.1 wrapper tool for Cutadapt (Martin, 2011) was used to remove adaptor contamination. High quality reads were mapped to mouse mm10 reference genome using Bowtie2 v2.2.9 (Langmead et al., 2009). After alignment, PCR duplicates were removed using picard v2.9.2 (Picard2019toolkit, 2019) and orphan reads were also discarded. Finally, uniquely mapped reads in proper pair were extracted using SAMtools v1.7 (Li et al., 2009). To perform peak calling on each replicate (n=2 biological replicates), MACS2 v2.1.1 (Zhang et al., 2008) was used with settings -g mm -p 1e-3 --nomodel --extsize --call-summits -B --SPMR. The value of --extsize was set based on MACS2 predicted fragment length. To obtain coverage tracks per replicates, --SPMR option of MACS2 was used to generate signal file of fragment pileup per million reads. Evidence from biological replicates were combined using MSPC v2 with parameters -r biological -s 1E-5 -W 1E-2. The resultant peaks were assigned the lowest p-value of the overlapping peaks across both replicates. Peaks corresponding to X, Y and M chromosomes were discarded and only reproducible peaks from both replicates were used as consensus peak set (Jalili et al., 2018). This resulted in identification of 11’422 regions significantly enriched in TBX3^3xF^ chromatin complexes that were used for further analysis. The raw sequencing data of the HAND2^3XF^ ChIP-seq from mouse limb buds at E10.5 (GSE55707, Osterwalder et al., 2014) were subjected to quality check using FastQC v0.11.4 (Andrews, 2010) and Trim_Galore v0.4.1 wrapper tool for Cutadapt (Martin, 2011). High quality reads were mapped to the mouse mm10 genome assembly using Bowtie2 v2.2.9 (Langmead and Salzberg, 2012). After alignment, PCR duplicates and orphan reads were discarded using SAMtools v1.7 (Li et al., 2009) and only uniquely mapped reads retained. Subsequently, MACS2 v2.1.1 (Zhang et al., 2008) was used with settings -g mm -p 1e-3 --nomodel --extsize 200 --call-summits -B –SPMR to perform peak calling. Peaks corresponding to X, Y and M chromosomes were discarded, which yielded 19’639 genomic regions significantly enriched in HAND2^3xF^ chromatin complexes.

### Annotation and evolutionary conservation analysis

The significantly enriched TBX3^3xF^ ChIP-seq peaks were annotated to promoter, intergenic and intragenic regions out using the annotatePeaks.pl utility available as part of HOMER (Heinz et al., 2010) and their evolutionary conservation was determined using the Phastcons conservation scores. The genome-wide track (track name: mm10.60way.phastCons60wayPlacental.bw) of the base-pair Phastcons scores in placental mammals was downloaded from the UCSC genome browser (Tyner et al., 2017). The base-pair scores for the 300bp flanking regions on either side of the TBX3^3xF^ peaks was determined using bwtool (Pohl and Beato, 2014).

### Motif enrichment analysis

To scan for consensus transcription-factor binding sites and de novo motifs in the 3057 TBX3 ChIP-seq peaks that are located in open chromatin regions (i.e. overlapping ATAC-seq peaks, Table S2) the *findMotifsGenome.pl* utility of the HOMER suite (v4.11, Heinz et al., 2010) was used with the following parameters: *size* of the region used to identify motifs was defined as number of bases downstream and upstream of the centre of the called ChIP-seq peak: -150,150. The motif length *len* was defined as the range from 6-18 bases. HOMER randomly selects background sequences from a pool of gene-proximal regions (+/- 50kb) for ChIP-Seq analysis and automatically discards genomic regions overlapping with the target peaks. These parameters were used to scan all TBX3 ChIP-seq peaks identified.

### Differential gene expression analysis using RNA-seq

polyA^+^RNAs were prepared from pairs of forelimb buds isolated from wildtype and *Tbx3*-deficient mouse embryos at E9.75-10.0 (28-31 somites). Similarly, polyA^+^RNAs were extracted from pairs of forelimb buds of wildtype and *Hand2*-deficient limb buds at E10.0-10.25 (31-33 somites). For each genotype, 3 independent biological replicates were analysed by RNA-seq using the Illumina NextSeq 500 to generate single-end reads of 75bp length. Raw sequencing reads were subjected to quality check using FastQC v0.11.4 and high-quality reads were aligned to the mouse (mm10) genome using STAR v2.5.2 (Dobin et al., 2013) aligner with --*twopassMode Basic* and --*quantMode TranscriptomeSAM* settings. Subsequently, the *rsem-calculate-expression* utility of RSEM v1.3.0 (Li and Dewey, 2011) was used to quantify gene expression and transcript levels across all samples. The mouse reference gene annotation in GTF format was obtained from ENSEMBL (release 91). Prior to identification of DEGs, small-ncRNAs (snoRNA, miRNA, miscRNA, scRNA, and scaRNA) were filtered out. To further reduce noise and increase robustness, only genes with counts per million reads mapped (CPM) ≥1 per replicate were maintained. edgeR (Robinson et al., 2010) was used to identify differentially expressed genes by wildtype and *Tbx3* and *Hand2* deficient forelimb buds. Sequencing libraries were normalized using TMM normalization and differential expression between pairs of conditions were evaluated using excatTest utility of edgeR. False discovery rates were estimated using the Benjamini-Hochberg correction (Benjamini and Hochberg, 1995). DEGs with an absolute Fold Change (FC) cutoff of >1.2 and an adjusted p-value ≤0.05 were considered as significantly different and included in downstream analysis. Functional enrichment of the biological processes for differentially expressed genes was assessed by [Gene Ontology analysis](http://geneontology.org/).

### Identification of TBX3 and HAND2 target genes

The TBX3^3xF^ (n=3057) and HAND2^3xF^ (n=19639) ChIP-seq peaks located in regions of accessible/open chromatin, as determined by ATAC-seq analysis (Jhanwar et al., 2021), were associated to their candidate target genes by GREAT analysis ([Genomic Regions Enrichment of Annotations Tool](http://great.stanford.edu/public/html/), McLean et al., 2010). GREAT analysis was conducted using the mouse mm10 reference genome with advanced options and the rule that genes are located within ≤1MB of a ChIP-seq peak in accessible chromatin. Among these genes, the ones expressed differentially (DEGs) and associated with TBX3^3xF^ and/or HAND2^3xF^ ChIP-seq peaks were defined as candidate TBX3, HAND2 or common target genes in mouse limb buds.

### Gene Ontology analysis of differentially expressed genes and putative gene targets of TBX3 and HAND2

[Gene Ontology analysis](https://geneontology.org/) (release 2021-08-18; Gene Ontology Consortium, 2021; Mi et al., 2019) was done for following data sets: 1. up- and down-regulated DEGs identified by pairwise comparison of wildtype vs *Tbx3*-deficient and wildtype vs *Hand2*-deficient limb bud samples; 2. candidate TBX3 and HAND2 target genes. For each set, the biological processes with adjusted p-values ≤ 0.05 were consider as significantly enriched and the top 20 processes are shown in Figs S2 and S4.

### Hierarchical clustering, plots and statistical testing

Clustering, plots, and statistics were handled in the statistical computing environment R v3. The GO enrichment analysis plots with the top 20 enriched biological processes (FDR≤0.05) were generated using [Gene Ontology analysis](https://geneontology.org/). Heatmaps of DEGs and differentially expressed target genes were prepared using [Python v3.7](https://geneontology.org/).

### Immunofluorescence analysis

#### Using frozen sections

Embryos were collected in ice-cold PBS and fixed for 2 hrs at 4°C in 4% PFA/PBS. After washing with PBS, samples were then cryoprotected using a gradient of sucrose: 10% sucrose/PBS (w/v), 20% sucrose/PBS, 30% sucrose/PBS (1hr each) at 4°C. Embryos were then embedded 50:50 (v/v) OCT/30% sucrose. For immunofluorescent staining, 10µm sections were prepared. *Tbx3*^3xF/3xF^ or wildtype sections were washed 3×5min in PBS, once 30min in PBT and again 5min in PBS. They were blocked in 1% BSA in PBT for 1hr at RT and incubated overnight at 4°C with the monoclonal mouse anti-FLAG M2 antibody (Sigma, F1804) diluted 1:500 in 1% BSA/PBS. Sections were washed 3×5min in PBS, once in PBT and were incubated in the dark for 60min at RT with the goat anti-mouse Alexa 488 secondary antibody (Invitrogen, A-11007) diluted 1:500 in 1% BSA/PBS. Sections were finally washed 3×10min PBS, once in PBT (5min), nuclei counterstained in 1µg/mL Hoechst-33258/PBS (5min) and rinsed again 3×5min in PBS. Then they were mounted in Mowiol 4-88 and dried overnight at RT in the dark.

#### Whole mount immunofluorescence analysis

After fixation in 4% PFA/PBS for 24 hrs, embryos were kept in storage buffer (PBS with 0.005% Na-azide. To start the experiment, embryos were washed with PBS (3xtimes 5min) and permeabilized in 0.5% Triton-X100/PBS for 60 min with shaking at room temperature. Then the samples were blocked in 1%BSA,0.1%Tween-20 with 5% serum in 0.5% Triton-X100 in PBS for 45 min. Then the samples were incubated with primary rabbit polyclonal anti-TBX3 antibodies (1:200, ab99302 ABCAM) in blocking buffer for 3 days at 4°C with gentle shaking. After 6 washes in 0.5% Triton-X-00 in PBS the samples were incubated with donkey anti-rabbit secondary antibodies (1:1000, Alexa 647, A311573, Life Science Technologies) in 1%BSA, 5% serum and 0.5% Triton for 2 days 4°C with gentle shaking. After 6 washes in 0.5% Triton-X100 in PBS, the samples were incubated and then the embryos were counterstained in DAPI (1:1000) 0.5% Triton-X100 in PBS overnight at 4°C. Finally, the samples were mounted and images acquired as described below.

### Whole Mount *in situ* hybridization (WISH)

Per genotype and gene n≥3 independent biological samples are analysed. Embryos are collected in cold PBS and then fixed for 24hrs in 4% PFA. Then they are gradually dehydrated into 100% methanol and stored at -20°C. To start the experiment, pool marked wildtype and mutant embryos with matched limb buds in one tube per probe to allow for direct comparison of results. Next, the embryos are rehydrated and washed twice in PBT. After bleaching in 6% H_2_O_2_-PBT for 15 min at room temperature (RT), they are washed in PBT (5min each) and treated with proteinase K (10µg/ml) for 10 (E10.0) to 15min (E10.5). The proteinase K digestion is inactivated in 2mg/ml Glycine-PBS for 5 min at RT. Then embryos are washed twice with PBT for 5 min and then re-fixed in 4% paraformaldehyde- 0.2% glutaraldehyde in PBT for 20min. After two washes in PBT at RT, the embryos are pre-blocked to reduce background in prehybridization buffer for minimally 60min at 70°C. 10 µl of riboprobe for the gene of interest is diluted in 1ml of prehybridization buffer per tube, denatured at 85°C for 5 min, quenched on ice and added to the samples after the removing the prehybridization solution and hybridised overnight at 70°C.

The next day, samples are washed in 800µl prehybridization buffer for 5 min at 70°C, then each 5 min 400µl of 2xSSC buffer are added (3 times) at 70°C. This is followed by two washes (30min each) in 2xSSC,0.1% CHAPS at 70°C. An RNase treatment is done for 45min in 2xSCC,0.1% CHAPS and 20µl/ml RNase A. Then samples are washed twice in maleic acid (pH5.2) for 10min at RT, then 2×30min with maleic acid at 70°C. This is followed by 3×5min washes with TBS-Tween 0.1% (TBST) and pre-blocking in 10% w/v lamb serum in TBST for 1hr at RT. The blocking solution is exchanged with anti-Digoxigenin-AP Fab fragments (Sigma, 11093274910) diluted 1:2000 in 1% w/v lamb serum in TBST and incubated overnight at 4°C.

After extensive washes in TBST (3×5min and then 8x 30 min), the samples are equilibrated for 3×10min in NTMT buffer (2ml 5M NaCl, 5ml Tris-HCl pH9.5, 5ml 1M MgCl_2_, 1ml Tween-20 in 100ml H_2_O). The signal is developed in BM Purple (Roche 11442074001) in the dark and the signal checked regularly until the best possible signal-to-noise ratio is achieved. Weak signals can be developed overnight at 4°C. The development is stopped by 5×5min washes in PBT for 5 min and then the PBT is replaced by PBS. For long-term storage samples are stored in 2% PFA in PBS 4°C. Images are acquired using a Nikon SMZ25 stereo microscope with DS-Ri2 camera and the NIS-Elements BR5.11.00 software or a Leica MZ FL2 stereomicroscope with the Leica Application Suite X software.

### HCR^TM^ RNA-FISH

The mouse HCR^TM^ probe sets for the different mouse genes analysed and *LacZ* mRNAs together with amplifiers and buffers were purchased from Molecular Instruments (USA). Briefly, embryos were fixed in freshly prepared 4% paraformaldehyde overnight at 4°C and dehydrated into 100% methanol for storage at -20°C. The HCR^TM^ analysis was done exactly as described in the recently published step-by step protocol by (Morabito et al., 2023) with the following small modifications: the photochemical bleaching was done for only 1 hour and the clearing before embedding was done overnight using refractive index matching solution (RIMS) with a RI of 1.45 for confocal imaging.

### Fluorescent Image Analysis

Immunofluorescent images of sections were acquired using the Leica SP5 confocal microscope and software, and processed using ImageJ software. Whole limb buds (immunofluorescence and HCR^TM^) were imaged using a confocal spinning disc (Nikon Ti-E, Hamamtsu Flash 4.0 V2 CMOS camera). The image acquisition software VisiView Premier was used. The different conditions of an experiment were acquired with the same settings as they were all in the same tube. To compare pattern changes, the pixel range (min - max) for a given gene was the same across all genotypes within an experiment and the same embryonic stage. Background or low signal was removed for pixel values ≤250 for all genes analyzed, with exception of *Gli3* where the threshold ≤500 pixels. When all three signals had to be displayed, we used the classic red/green/blue combination, which is the only one that allows the different color combinations when working with three different overlapping signals.

### Mutations of the TBX3 and HAND2 motifs in the *Gli3* mouse core enhancers *mm1179* and *mm-hs1586* and generation of mouse *LacZ* founder embryos

The TBX3 (this study), HAND2 (Osterwalder et al., 2014), HOXD13 (Sheth et al., 2016), SMAD4 (Gamart et al., 2021) and ß-Catenin ChIP seq-profiles (Sheth et al, unpublished) were used together with the H3K27ac profile were used to map the core regions in both enhancers. The candidate HAND2 binding sites were mapped using the HAND1::TCF3 motif (Jaspar ID MA0092.1) and the TBX3 binding sites using the human TBX3 motif (Jaspar ID MA1566.1) with ap-value cut-off of 0.01. Two of the most conserved bases of each motif were mutated (Fig. S7C), and the mutated binding regions rescanned to verify their inactivation as HAND and TBX binding regions, respectively. The mutated core enhancer regions (Fig. S8) were then synthesized (IDT, Integrated DNA Technologies), inserted into the *Hsp68*-lacZ vector (Pennacchio et al., 2006) and founder embryos generated by pronuclear linear DNA injection.

### Generation of mouse *LacZ* reporter strains for two *Gli3* enhancers

The enhancer constructs were generated using a PCR based strategy from mouse genomic DNA. Primers were designed using Primer3, software. The amplified DNA was inserted into the *Hsp68*-lacZ vector using the Gibson Assembly kit (E2611S, New England Biolabs). The resulting constructs were verified, injection grade DNA produced and linearized. The linearized plasmids were for pronuclear injection using the University of Basel Centre for transgenic mice (CTM) and all founder mice genotyped using specific primers. Two founder mice were obtained for the *Gli3 mm1179* and three for the *mm-hs1586* enhancer. Initial analysis to verify their known enhancer activities was done before crossing one line each into our *Tbx3^flox^* line to comparatively analyse the enhancer activities in wildtype and in *Tbx3* mutant mouse embryos. All primers are listed in Table S10.

### Analysis of transgenic *LacZ* embryos

It is standard to determine tissue-specific enhancer activity is n≥3 independent transgenic embryos expressing the *LacZ* reporter in the tissue of interest (Visel et al., 2007). Embryos are collected in ice-cold PBS and then fixed in PBS with 1% formaldehyde, 0.2% glutaraldehyde, 0,2% NP40, 0,01 sodium deoxycholate in PBS) for 20 min at 4°C. After 3 washes in PBS at room temperature, the embryos are stained at 37°C in the dark with gentle rotation. The staining solution consists of 0,5mg/ml 5-Bromo-4-chloro-3-indolyl β-D-galactopyranoside (X-Gal) in dimethyl formamide, 0,25mM of K3Fe(CN6), 0.25mM of K4Fe(CN6), 0.01% NP40 and 4mM of magnesium chloride. Embryos are checked for the blue staining due to ß-Galactosidase activity after about 90 mins and then every 40-60 min. If the staining is weak or absent, the staining solution is replaced and the development is continued overnight at 37°C in the dark. The next day, embryos are washed in PBS and post-fixed in 4% PFA/0.1% glutaraldehyde in PBS for 1 hour: washed twice in PBS and then stored in PBS with 0,05% Na Azide at 4°C. Samples are analysed and z-stack images acquired using a Nikon SMZ25 stereomicroscope equipped with a DS-Ri2 camera and the NIS-Elements BR5.11.00 software.

### Available datasets used in this study

A list of genes expressed in mouse limb buds and/or whose mutagenesis, loss-of-function, and gain-of-function causes congenital limb malformations based on information was manually curated using information available via the *[embrys](https://www.embrys.jp/embrys/2DView/pictures)* and the [Mouse Genome Informatics (MGI) databases](http://www.informatics.jax.org/). The raw sequencing ATAC-seq data from mouse forelimb buds at E9.75 corresponds to the dataset published by (Jhanwar et al., 2021) (GSE164736). The HAND2 ChIP-seq dataset from mouse limb buds at E10.5 is from Osterwalder et al., 2014 (GSE55707). The histone modification ChIP-seq datasets (H3K27me3 and H3K27ac) for mouse limb buds at E10.5 are from the mouse [ENCODE project](https://www.encodeproject.org/data/).

## Supporting information

Supplementary Figures and Inventory

## Data availability

Newly generated genome-wide dataset of TBX3ChIP-seq and *Tbx3* and *Hand2* RNA-seq datasets are available through the Gene Expression Omnibus (GEO) repository under the series GSE192486.

## Acknowledgements

We are very grateful to V. Christoffels for making the mouse *Tbx3*^ΔVenus^ allele and to A. Moon and C. Cantu for making the *Tbx3*^flox^ allele available to us for the purpose of this study. We thank A. Offinger and her team for excellent animal care and welfare, A. Baur, J. Stolte and O. Romashkina for expert technical assistance during the initial phase of the project and A. Morabito for help with Western blot analysis. P. Pelczar and his team (Centre for Transgenic Models at the University of Basel) for generating the *LacZ* founder embryos. We thank [enGene Statistics](https://www.engene.ch/) for the TBX3^3xF^ ChIP-seq peak distribution diagram and motif enrichment analysis on a fee for service basis. We are indebted to P. Lorentz and L. Sauteur from the DBM Microscopy Core Facility and Nikon Center of Excellence for training and advise in using the Visitron spinning disk confocal microscope and with image analysis.

## Author contributions

GS performed most of the in-depth analysis of the *Tbx3* (WISH) and shared *Hand2*-*Tbx3* target genes (HCR^TM^) plus all the *Gli3* enhancer analysis. This analysis resulted in all the major conclusions for the study, in particular the essential and unique role of *Tbx3* in posterior gene expression boundary formation and its role in up-regulating posterior genes together with *Hand2*. AG generated the *Tbx3*^3xF^ allele together with AZ and the CTM mouse core facility of University of Basel. AG characterised the *Tbx3*^3xF^ line, generated all genome-wide datasets for differential gene expression analysis from wildtype and mutant forelimb buds and performed the initial analysis of *Tbx3* target genes. SJ performed all bioinformatics analysis of the primary datasets and generated most of the bioinformatics data shown in the manuscript. VP generated definitive heatmaps for the *Tbx3* target GRN and the analysis of the shared *Tbx3*-*Hand2* target genes. RS generated the *Tbx*^3xF^ ChIP-seq datasets from early forelimb buds with help from AG and co-supervised the entire study. RZ and AZ manually curated the list of TBX3 target genes with known functions and spatial expression patterns to construct the scheme of the TBX3 target GRN in early limb buds. RZ and AZ conceived and supervised the study, acquired all funding and wrote the manuscript together with GS and input from all authors.

## Supplementary information

Supplementary Information includes Figures, Tables and Movies.

