## Supplementary Figures and Inventory for "TBX3 is essential to establish the posterior boundary of anterior genes and up-regulate posterior genes with HAND2 during onset of limb bud development"

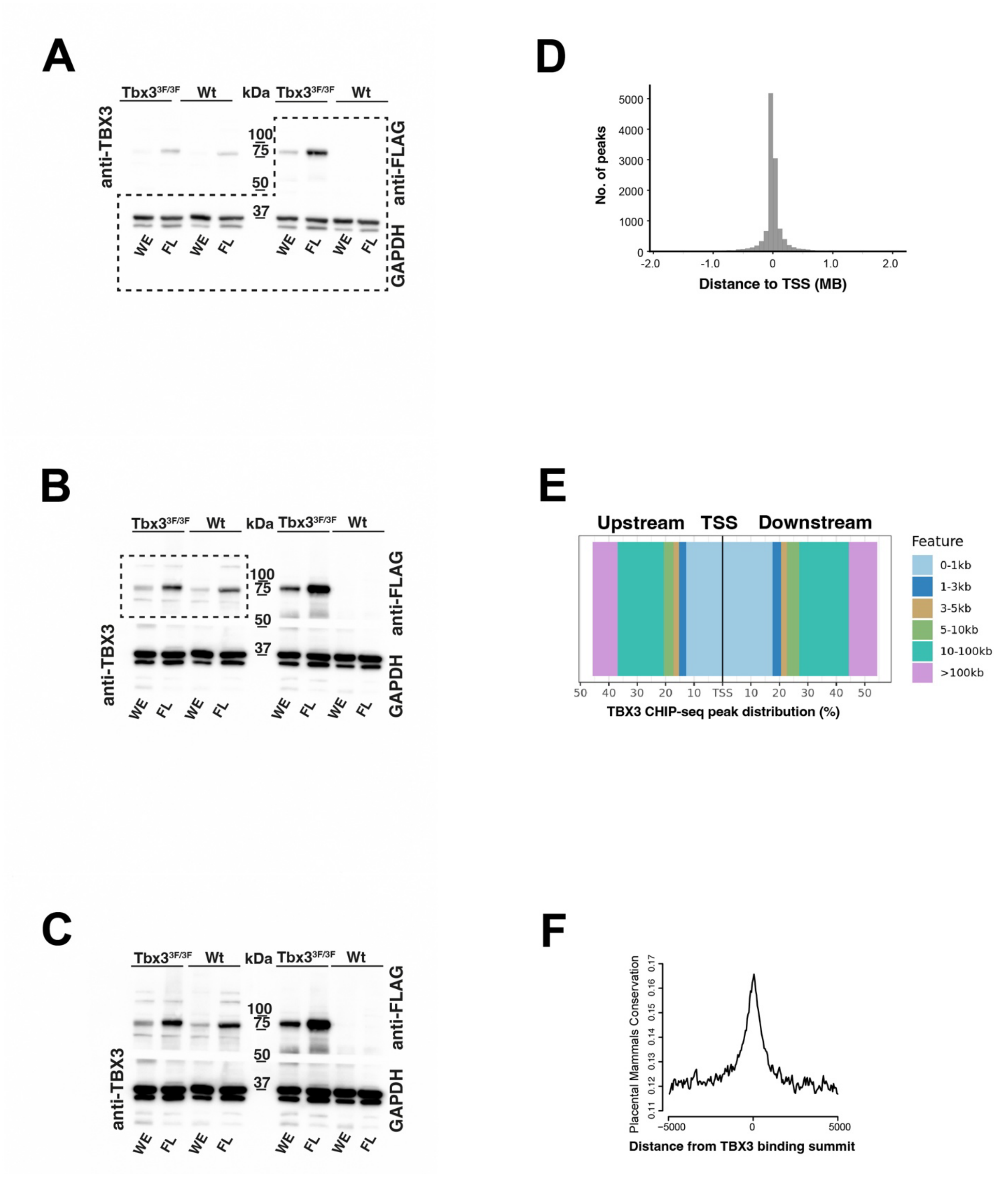


**Figure S1. Identification of the TBX3 cistrome using a novel mouse *Tbx3*$^{3xFLAG}$ allele.** (A-C) Different exposures of the uncropped Western blot used for Fig. 1B (exposure times are: 4 sec (panel A), 30 sec (panel B) and 90 sec (panel b). The M2 anti-FLAG specifically detects the mouse TBX3$^{3xFLAG}$ protein, but not the wild-type TBX3 protein (n=3). The dotted line outlines the cropped areas used for the composite shown in Fig. 1B. (D) Histogram shows distance distribution of the verified genomic regions enriched in TBX3 chromatin complexes with respect to the nearest transcriptional start site (TSS) at E9.75-E10.25 (28-32 somites). (E) Plot showing the fraction (%) of the $TBX3^{3xF}$ ChIP-seq peak distributions in relation to their distance from TSS. (F) Conservation analysis of the genomic regions enriched in TBX3 ChIP-seq using Phastcons conservation scores. Shown is the average Phastcons conservation of the TBX3-bound genomic regions (n=11,422) identified by ChIP-seq analysis.

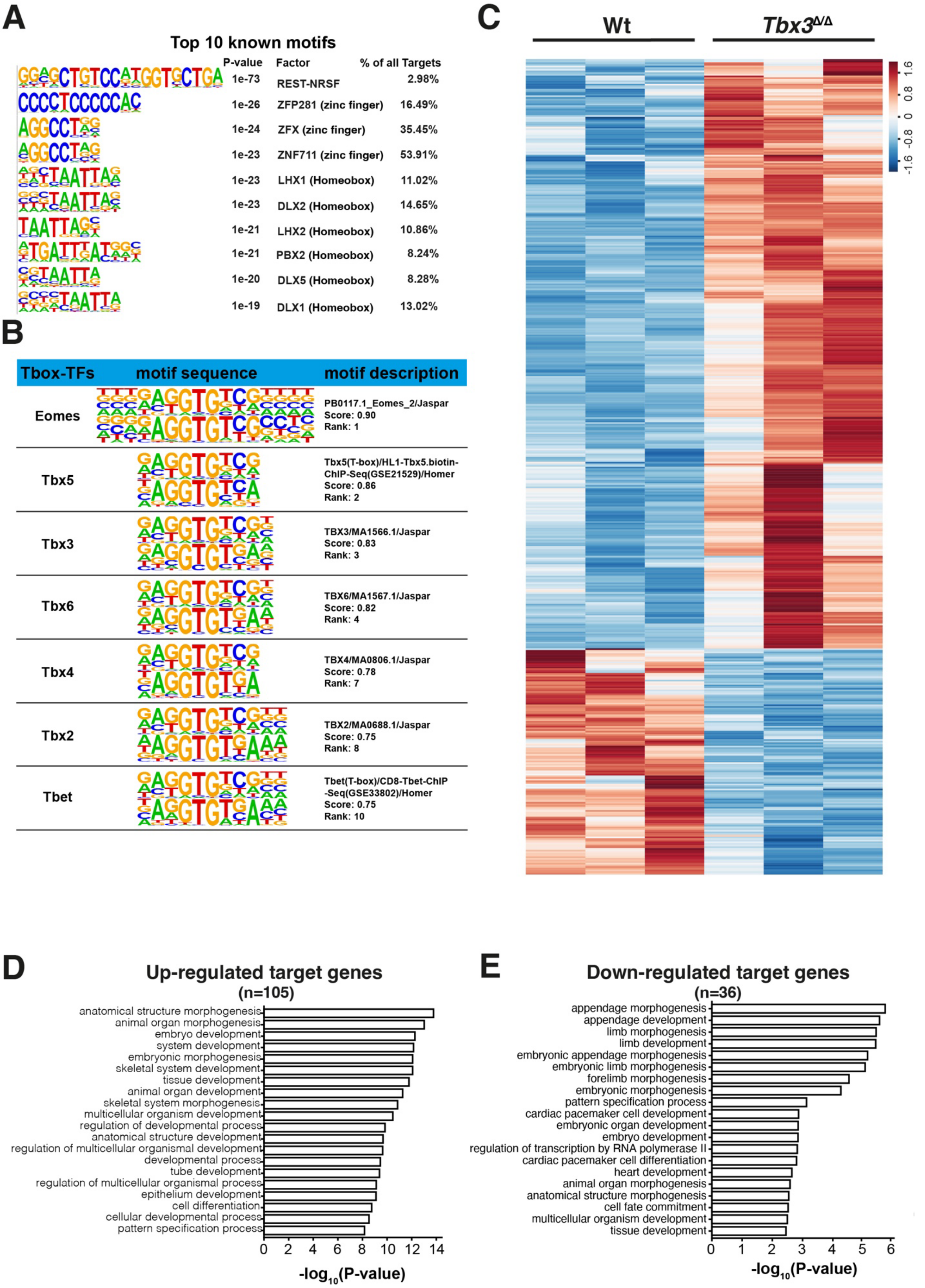
A
Top 10 known motifs
P-value
Factor
% of all Targets
1e-73 REST-NRSF 2.98%
1e-26 ZFP281 (zinc finger) 16.49%
1e-24 ZFX (zinc finger) 35.45%
1e-23 ZNF711 (zinc finger) 53.91%
1e-23 LHX1 (Homeobox) 11.02%
1e-23 DLX2 (Homeobox) 14.65%
1e-21 LHX2 (Homeobox) 10.86%
1e-21 PBX2 (Homeobox) 8.24%
1e-20 DLX5 (Homeobox) 8.28%
1e-19 DLX1 (Homeobox) 13.02%
B
Tbox-TFs
motif sequence
motif description
Eomes
PB0117.1_Eomes_2/Jaspar
Score: 0.90
Rank: 1
Tbx5
Tbx5(T-box)/HL1-Tbx5.biotin-ChIP-Seq(GSE21529)/Homer
Score: 0.86
Rank: 2
Tbx3
TBX3/MA1566.1/Jaspar
Score: 0.83
Rank: 3
Tbx6
TBX6/MA1567.1/Jaspar
Score: 0.82
Rank: 4
Tbx4
TBX4/MA0806.1/Jaspar
Score: 0.78
Rank: 7
Tbx2
TBX2/MA0688.1/Jaspar
Score: 0.75
Rank: 8
Tbet
Tbet(T-box)/CD8-Tbet-ChIP-Seq(GSE33802)/Homer
Score: 0.75
Rank: 10
C
Wt
Tbx3Δ/Δ
1.6
0.8
0
-0.8
-1.6
D
Up-regulated target genes
(n=105)
anatomical structure morphogenesis
animal organ morphogenesis
embryo development
system development
embryonic morphogenesis
skeletal system development
tissue development
animal organ development
skeletal system morphogenesis
multicellular organism development
regulation of developmental process
anatomical structure development
regulation of multicellular organismal development
developmental process
tube development
regulation of multicellular organismal process
epithelium development
cell differentiation
cellular developmental process
pattern specification process
0 2 4 6 8 10 12 14
-log10(P-value)
E
Down-regulated target genes
(n=36)
appendage morphogenesis
appendage development
limb morphogenesis
limb development
embryonic appendage morphogenesis
embryonic limb morphogenesis
forelimb morphogenesis
embryonic morphogenesis
pattern specification process
cardiac pacemaker cell development
embryonic organ development
embryo development
regulation of transcription by RNA polymerase II
cardiac pacemaker cell differentiation
heart development
animal organ morphogenesis
anatomical structure morphogenesis
cell fate commitment
multicellular organism development
tissue development
0 1 2 3 4 5 6
-log10(P-value)

**Figure S2. TBX3 binding motif analysis and differentially expressed genes in *Tbx3* deficient limb buds (E9.75-10.0)** (A) Top known motif analysis of genomic regions enriched in TBX3 chromatin complexes and accessible in mouse embryos forelimb buds at E9.75-10.25. (B) HOMER de novo motif analysis reveals the high score ($x > 0.7$) T-box motifs enriched by ChIP-seq analysis sand shows the similarities of the binding motif sequences identified for different TBX transcription factors. (C) Heatmap of all DEGs (n = 494) at E9.75-10.0 (28-31 somites) illustrating their relative gene expression ratios acrosss wild-type and *Tbx3*-deficient forelimb buds (n=3 biological replicates). Only DEGs with an absolute fold-change (FC) cutoff of ≥1.2 and an adjusted $p$ value ≤ 0.05 were considered significantly changed. Among these 494 DEGs, 357 are up-regulated and 137 DEGS are down-regulated. The z-score scale represents mean-subtracted regularized log-transformed read counts. (D)Top enriched (n=20) biological processes identified by gene ontology (GO) analysis for the up-regulated TBX3 target genes (n=105). (E) Top enriched (n=20) biological processes identified by GO analysis for the down-regulated TBX3 target genes (n=36). DEG: differentially expressed gene.

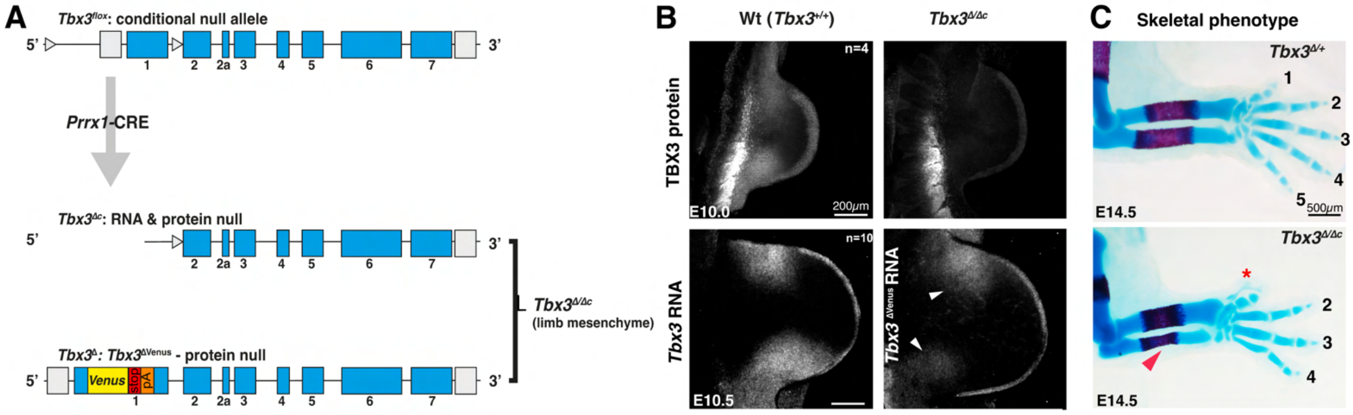


**Figure S3. The different *Tbx3* alleles used for analysis.** (A) Scheme illustrating the generation of *Tbx3*$^{\Delta/\Delta c}$ mouse embryos, in which *Tbx3* is conditionally inactivated in the limb bud mesenchyme using a *Prxx1-Cre* driver. The LoxP sites are indicated by open arrow heads. Conditionally deleting one *Tbx3*$^{flox}$ in the context of the *Tbx3*$^{\Delta c}$ allele results in rapid clearance of the TBX3 protein (panel B). (B) Top panels immunofluorescence analysis show the clearance of TBX3 proteins from *Tbx3*$^{\Delta/\Delta c}$ forelimb buds by E10.0 (29 – 33 somites, n=4). Bottom panels: HCR$^{TM}$ detection of Tbx3 mRNAs in wild-type and *Tbx3*$^{\Delta/\Delta c}$ forelimb buds (n≥4). The HCR$^{TM}$ probe set detects the *Tbx3* protein-null transcript in *Tbx3*$^{\Delta/\Delta c}$ forelimb buds at variable levels (indicated by white arrow heads, E10.5, 34-36 somites). White arrows indicate the remaining *Tbx3* mRNA signal. Scale bar: 200$\mu$m. (C) Skeletal morphology of *Tbx3*$^{\Delta/\Delta c}$ forelimbs at E14.5 shows the classical *Tbx3* mutant limb skeletal phenotype. The asterisk indicates the duplication of digit 1 and digit 5 is lost (shown here) or hypoplastic. The red arrow points to the hypoplastic ulna. Scale bar: 500$\mu$m.

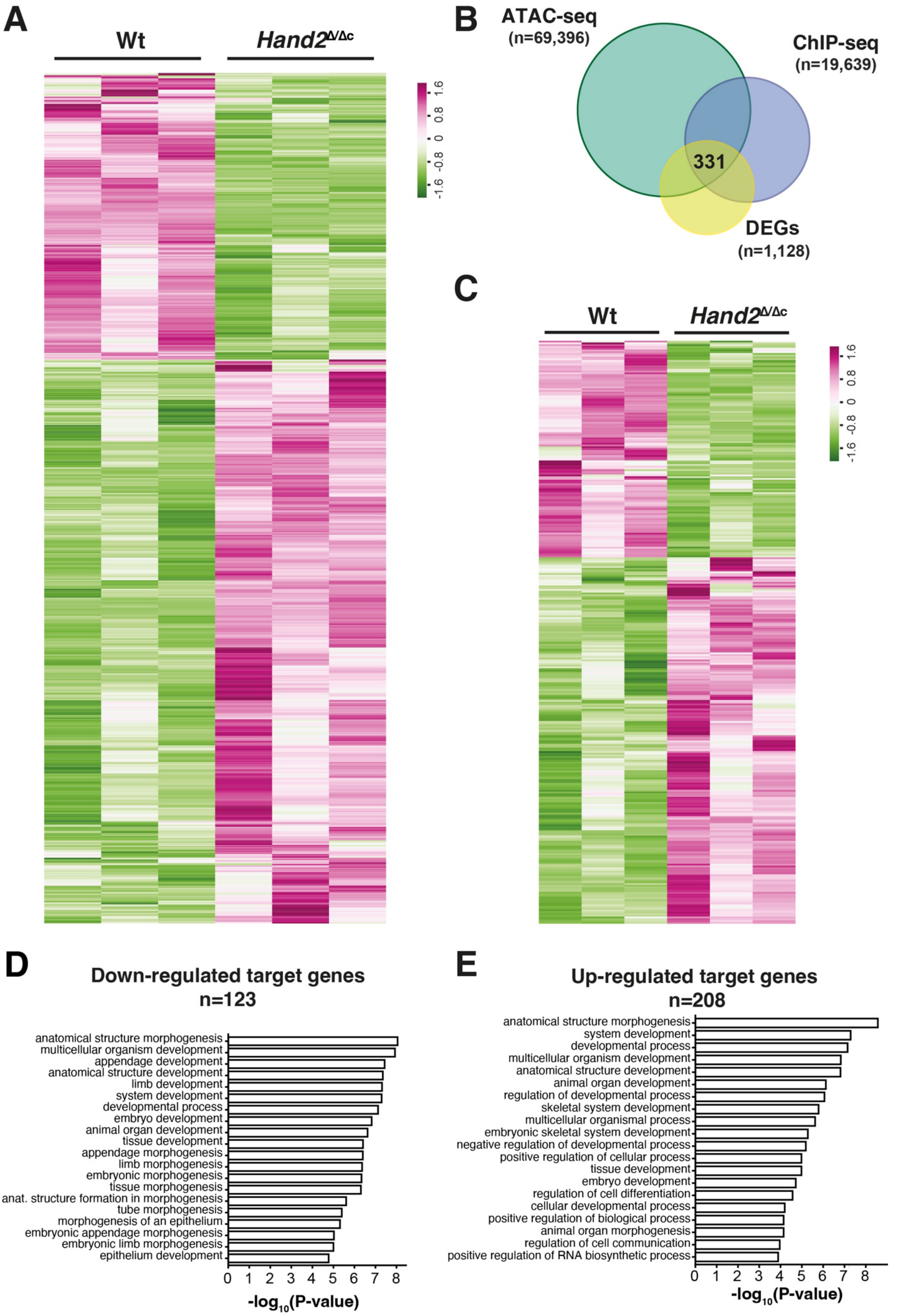
A
Wt
Hand2Δ/Δc
1.6
0.8
0
-0.8
-1.6
B
ATAC-seq
(n=69,396)
ChIP-seq
(n=19,639)
331
DEGs
(n=1,128)
C
Wt
Hand2Δ/Δc
1.6
0.8
0
-0.8
-1.6
D
Down-regulated target genes
n=123
anatomical structure morphogenesis
multicellular organism development
appendage development
anatomical structure development
limb development
system development
developmental process
embryo development
animal organ development
tissue development
appendage morphogenesis
limb morphogenesis
embryonic morphogenesis
tissue morphogenesis
anat. structure formation in morphogenesis
tube morphogenesis
morphogenesis of an epithelium
embryonic appendage morphogenesis
embryonic limb morphogenesis
epithelium development
0 1 2 3 4 5 6 7 8
-log10(P-value)
E
Up-regulated target genes
n=208
anatomical structure morphogenesis
system development
developmental process
multicellular organism development
anatomical structure development
animal organ development
regulation of developmental process
skeletal system development
multicellular organismal process
embryonic skeletal system development
negative regulation of developmental process
positive regulation of cellular process
tissue development
embryo development
regulation of cell differentiation
cellular developmental process
positive regulation of biological process
animal organ morphogenesis
regulation of cell communication
positive regulation of RNA biosynthetic process
0 1 2 3 4 5 6 7 8
-log10(P-value)

**Figure S4. Identification of differentially expressed HAND2 target genes in mouse forelimb buds.** (A) Heatmap of DEGs (n = 1128; down-regulated: 380, up-regulated: n=748) in wild-type and *Hand2*$^{\Delta/\Delta c}$ forelimb buds (E10-10.25, 29-32 somites, n=3 biological replicates per genotypes). DEGs with significant changes in transcript levels must have an absolute FC cutoff of ≥1.2 and an adjusted *p* value ≤ 0.05. The z-score scale represents mean-subtracted regularized log-transformed read counts. (B) Shown is the intersection between the significantly enriched HAND2-bound regions (ChIP-seq, Osterwalder et al., 2014), open chromatin regions (ATAC-seq, E9.75) and differentially expressed genes (DEGs) between wild-type and *Hand2*-deficient samples (RNA-seq, E10-10.25). A total of 331 HAND2 candidate gene targets are identified in the mouse forelimb buds. (C) Heatmap illustrating the relative gene expression of candidate gene targets of HAND2 (n=331) that showed significant changes (absolute FC cutoff of ≥1.2 and an adjusted *p* value ≤ 0.05) between wild-type and *Hand2*-deficient samples during RNA-seq analysis. The z-score scale represents mean-subtracted regularized log-transformed read counts. (D) Top (n=20) enriched biological processes identified by GO analysis of up-regulated target genes (n=208) of HAND2. (E) Top (n=20) enriched biological processes identified by GO analysis of down-regulated target genes (n=123) of HAND2. DEG: differentially expressed genes, GO: gene ontology.

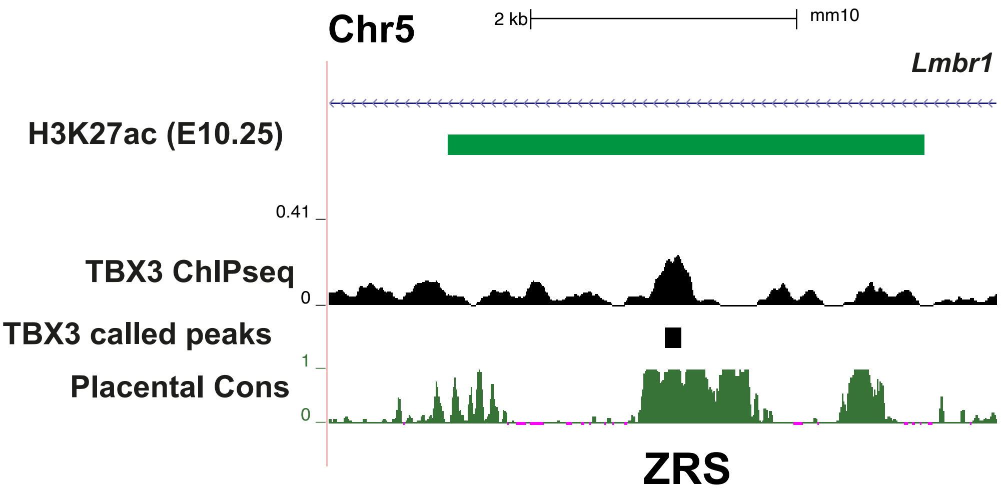


**Figure S5. A TBX3 ChIP-seq peak in the ZRS points to direct regulation of *Shh* expression in mouse forelimb buds.** UCSC browser view (5kb) of the ZRS *Shh* limb bud enhancer in the *Lmbr1* locus. H3K27ac modified region marking active chromatin is indicated by a green line with a green rectangle, the called TBX3 ChIP-seq peak is marked by a black square. The Placental cons shows the base pair conservation.

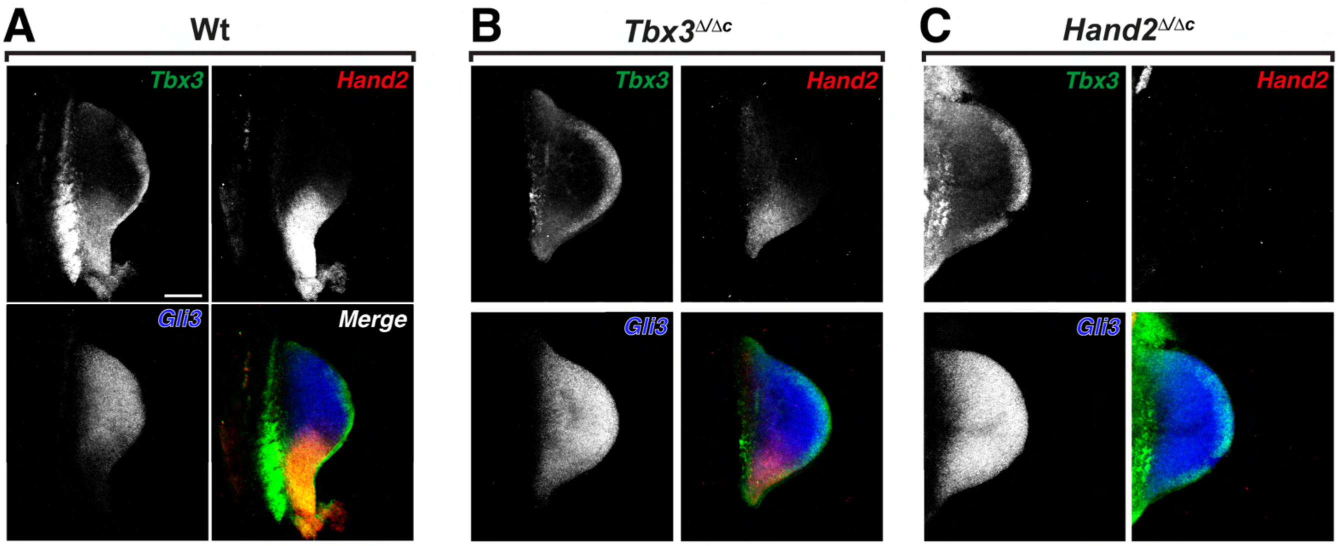


**Figure S6. *Tbx3* is required to restrict *Gli3* from the posterior mesenchyme and posterior *Tbx3* expression is lost in *Hand2*$^{\Delta/\Delta c}$ forelimb buds**. (A-C) HCR™ analysis of the *Tbx3*, *Hand2* and *Gli3* expression in wild-type (panel A), *Tbx3*$^{\Delta/\Delta c}$ (panel B) and *Hand2*$^{\Delta/\Delta c}$ forelimb buds (panel C) at E10.0 (29-32 somites). n=3. Scale bar: 200µm.

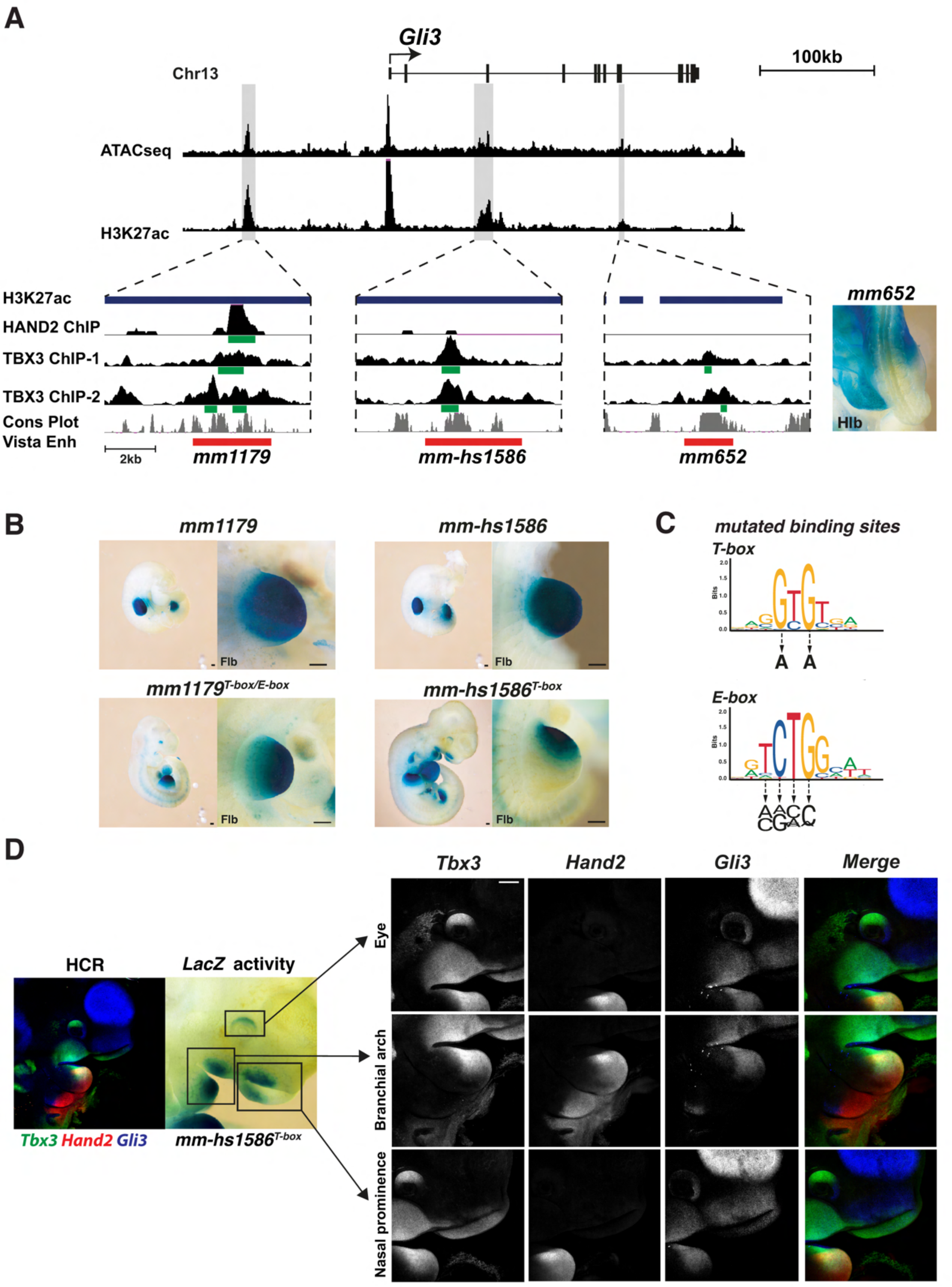


**Figure S7. TBX3 and HAND2 bind to *Gli3* enhancers and their motif are necessary for their correct spatial activity.** (A) UCSC browser view of the *Gli3* genomic landscape with enlargements (below) of the relevant regions of accessible and active chromatin (ATAC-seq and H3K27ac peaks). The two limb enhancers *mm1179* and *mm-hs1586* plus a third potential enhancer *mm652* are highlighted (grey shading). The enlargements below show of the genomic locations of the HAND2 and TBX3 ChIP-seq peaks mapping to these enhancers and the genomic regions tested for *Gli3* enhancer activity (red bars). The right-most panel shows the transient early activity of the *mm652 LacZ* reporter construct in hindlimb buds (Hlb) of a transgenic founder embryo at E10. 5 (n=5/6). (B) Analysis of the *LacZ* reporter activity of the wild-type (*mm1179*: n=3; *hs-mm1586:* n=7; top panels) and mutant *mm1179* (n= 6) and *hs-mm1586* (n=11/13) enhancers (with mutated *T-box* and *E-box* motifs, lower panels) in transgenic mouse founder embryos at E10.5 (34-36 somites). Enlargements show forelimb buds (Flb). Scale bar: 200μm. (C) Scheme showing the base mutations to inactivate in the *E-box* and *T-box* motifs in the core regions of both enhancers (*mm1179*: 19 *E-box* motifs and 18 *T-box* motifs; *mm-hs1586*: 15 T-box motifs). (D) Several regions in the developing head (eye, nasal prominence and branchial arch show ectopic *LacZ* expression of the *hs-mm1586* enhancer with mutated *T-box* motifs (n=4). All these regions express *Tbx3* and *Gli3*, while *Hand2* expression is only detected in the 1$^{st}$ branchial arch. Scale bar: 200μm.

# A

**mm1179_ fragment 1 -repeat sequence- fragment 2: mutations of T-box and E-box motifs**

CCCTCACTAAAGGGAACAAAAGCTGGTACCGAAGAGTGTTACCGGCCTCTAAGTGAGACCTGGGAA
ATTAGCCCTTTTGTTGAACTGATCTTAATTTCCTTTGGGTCACAAATGTCCCCTCAGGAAAACAGGAC
TCACAGCCCTGGCCAATAAAGACCACCCCACTAATTAATTGAGTCATCTATATCTTGCAAGTGCAAG
ATGAAGCATTTAAATCTAAATGTTTCATATTTTAAATAGAAACGGCCAAAGTCATGGGGCTTCAATA
TGAATGTATTCCCCGGTAAAAAAAAGTATCTACTAAAGAAGCTTCCCGGGCTCAGGCAGAGCTCC
GGGTCCTCTGAAAATCAAATTTAAATGGCCCCGAGCAGCGGAGAACATATCTCCTCAGAAAAACAAC
GGGGCCTTTGTCTAATTGCCCATTTATTACGGTTTTATCTCAGTCCCCCAAAGAAAAGGAAAAGAAG
GGGACTGATTTAGTCCTCTTCATCCACAACTTTAACGGGCATGTGGTTCTAATGAAGTCGAGACTTT
CGCCTAACGAAATGTAATATAAACTCCCCTAATTGAAAAGTAATAAAAACATCAAAAACAATATCAAT
TAGCCAGTGAAGGGCCCCTTGCCTCTTTTGAAGACGGCTAGTGGTCCCGGAAAGTACTCTAAGCTA
TACACAAGAGGACACACA

- repeat sequence (177bp)-

ACACCCCCATTTCCAAACGTTTCCTCGGGGAATGACAAGAAAGAGCATCCCTGGTTAATATCTAGCC
ACTAAATATATATTGAAATCTCTAACTTGGGCAAGATCCAAACATTCAATAAAAGAATGGGGGGCCG
GATACCGTCAAAATAATAACAAAGGAGCATATCCGTTAATTATGATCTGAGAGTCAATAAAGTCATCT
GATGGCCCATTTAAGGGAAATCATTAGAGAGTAGAATTTAGGGGCAAAGAACTGCAGGCTTCCTTC
CTCCAAAAGCTACCCATTTCATTAATATCTTCAAAAAGGATGAAGGCCCCGCCCGAGCCATTGCCCC
TAGAGAAGGGGCCTAAACCGGAACTTTCCATTTCTCATAATTGAGATCACGGGAATAATCAGCTCTC
TTTTTAAGAAACCAGCCAAGACTAAATGGCAGCAAGACTGCTGGAGGTGCAAAGTCACAGGCTTCAA
GAGTGATTTCCATGACCACATAGTGTGACTTGCTGATGGGGAACTAGGAAGAGCTGGGCTGTGAAA
GATGTCTGCAGGACCCATGCAGTTCATTTTCCTATGGATGGAATTGGTTTTTTGGGCTCTAGGATTC
AAGTCTCTTTAAGCAAGCAAAGCTGTAAACAGAATACGTTTACCTCTGCTTTCTAGCTGCCAAAACAA
CAGAGGAGAGGGGTACCGAGCTCCAGGAACATCCAAACTGA

# B

**mm-hs1586 mutations of T-box motifs**

GTGTGTGTGTGTGTAATAGTTTTGGGTATTCCAATGCTTAGTGATTTGTGGAAACAAACTAATTTTAA
GAATTCAATACTAAATTTGTTACTATGGAAGTTTTTATTGTGAGCCCAACTCTCCTTATGAGCCTGCC
AGAGACACTGGAAGTCGCCTTTCTAGTCCTAACTGAATTAAAGAAATTAAGCAGGCCTCTTGGAGTA
TAAGTCTAGGACAGTTCTTTTCAAAGGCAGCCTATAAGGTAAATCCATACGGGTTGAGTTTACCTCA
GGCCTGTTTCTGGTTTAAGAATATAATTTGGCTCTGGAGTAATGAAAGTCATGTTGAGATAGCAAAG
CTGGAAGCTCCTGATTACAGATTTATAAACTTGCCAGGAAATAGGCTCGCCATTTAACCTTCCTGCC
TCCAGTTCCAGATGATCCATTGAGCTCCAGAGAAAGAGACTGATATGTTTCCCTTTTCCAAAATCGT
TCAGAAATTTTAATATTTTAAGGCAAAAGGTCATTATCCAGATGACTGATTAGTGAGGACAGCTGATT
AGAGCATATTATTTTGCTCAAGAAAATACCCAGGATAAAAGCTAATTTCATCCCTTCATTAGTTGGCT
GCAAGCAAGGGGAAGGCTCACTTATTCCTCAGATCAAAGAGACTCGAAGCAACCTTAAGTTCCAGC
AACTGCTTCTCCTATCAGAATATGTTGGTGGAAAATATGGCTATGCAATAGTTTCTAAGTAGAAAGGA
CAGTTTTTGTCAGAAGCATGAAAATACGATTGTTTGGATTTTGGGGAGGGGCAGTGGTTGGCTTTTA
AAAAGTTAATTGCTTTTCTTGAAAAAAAAATTCCCATTGGTCCCATTATTGAAAAGATATCTGTTGAAG
CAGAAACAAAGACAGAGGGGTTTGGACAACTATTCTTCCTTCCTCCTTTCAGGCTTACTTCATTTTGT
TTTTTCTTGACTTTTAGGAAGGAAGATTTAAAATAATATTGGTAGGATGTTTTACTCTGGTCACATTTT
GAAGTCTTTAGTTATATTTTTATTTGTGGTCTCTTTTGAGTATCATTCATGCAGAAACAGTAGGGATTC
CTATAGAGTCCCCACCTGGGTTTATATATCCGCATGTTAAGTACACTCACTGTGCTGTTTAAGCTGC
CCACAAGCTCCTCCCAGTGTTTTGTATTTCTGTGTGAACACAAGCTTTTCCCATCACAGCGATGTTTT
GTTTTGTTTTTTCCAATGCCCACTAATTAGTGAGGTGTTTAATTAGTCCAGACAGCTGATGAGCACTT
CTTTCTATTGGCTGATGGTAGTGGAACCGTTAACTTTCTATCCAGTTTGCAGCTTGTTCTTGCGTCCT
TTGTTTTAACGGTTTCCCACCCAGGCGTCTCTCCATCCTTTGAGGGACTGGCTTTCCCGTCACTGGA
ACCCCTGTTCTCACCCAGTGGTCTGTGTGACCCATTTGCTGGTTCACCAAAATAATGTTAACGAACA
CTGCCTGGAGATCAAATATTATGGTTTAGACTTATTTATTGCTTTCTAGTTTGAAGATATTGTACTTGT
AAGTTGGTTATTCTTGTGATTTTCCTACGTTACTATACAAGTGAATGTAGAAGTAACCTTCCAGTCTT
GAGCTGGAGCCCCTCGGCAAGTCAGTGTTGAGTCCCCCACCCCCAAATGTTTTCCCTACCTTGGGG
ACATTGTGTAGATAGCTATGTGCAGAAAGCCTAGAAAACTGAACAGACTGGCGGTACCGAGCTCCA
GGAACATCCAAACTGA

**Figure S8. Sequence of the mm1179 and mm-hs1586 *Gli3* enhancers with the nucleotide mutations in T-box and E-box motifs.** (A) Mutated *mm1179* core regions with all nucleotide changes shown in bold. (B) Mutated *mm-hs1586* core region with all nucleotide changes shown in bold. T-box motifs are indicated in yellow and E-boxes are underlined.

**INVENTORY OF SUPPLEMENTARY TABLES**

**Table S1.** Annotation of curated TBX3 ChIP-seq peaks to nearest promoter and genomic features.

**Table S2.** Association of the curated TBX3 ChIP-seq peaks to the two nearest genes (range ≤1Mb).

**Table S3.** List of differentially expressed genes between wildtype and T*bx3*$^{\Delta/\Delta}$ limb buds at E9.75-10.0.

**Table S4.** List of differentially expressed TBX3 target genes.

**Table S5**. Manually curated functional annotation of TBX3 target genes. Grey shaded genes are expressed during limb bud development. Genes indicated in bold are transcription factors.

**Table S6.** TBX3 target genes that are also differentially expressed in wildtype versus *Shh*-deficient limb buds at E10.5.

**Table S7.** List of differentially expressed genes between wildtype and *Hand2*$^{\Delta/\Delta}$ limb buds at E9.75-10.0.

**Table S8.** List of differentially expressed HAND2 target genes.

**Table S9.** Shared TBX3 and HAND2 target genes.

**Table S10.** Primers for genotyping mouse strains

**SUPPLEMENTARY MOVIES 1 and 2**

**Movie 1.** HCR$^{TM}$ analysis of a wild-type mouse limb bud at E10.5 (35 somites) showing the spatial expression of endogenous *Tbx3* (red), *Gli3* (green) and the transgenic *mm-hs1586-LacZ* reporter (blue). **Movie 2.** HCR$^{TM}$ analysis of a *Tbx3*$^{\Delta/\Delta c}$ mouse limb bud at E10.5 showing the spatial expression *Tbx3* (red), *Gli3* (green) and the transgenic *mm-hs1586-LacZ* reporter (blue).

Limb buds are oriented with anterior to the top and posterior to the bottom, proximal to the left and distal to the right. The movies show scans through the z-stacks in ventral to dorsal direction. These scans through the entire limb bud mesenchyme show that the wildtype *Tbx3* expression does not significantly overlap the spatially identical *Gli3* and *LacZ* expression domains.
